# Locally Fixed Alleles: A method to localize gene drive to island populations

**DOI:** 10.1101/509364

**Authors:** Jaye Sudweeks, Brandon Hollingsworth, Dimitri V. Blondel, Karl J. Campbell, Sumit Dhole, John D. Eisemann, Owain Edwards, John Godwin, Gregg R. Howald, Kevin Oh, Antoinette J. Piaggio, Thomas A. A. Prowse, Joshua V. Ross, J. Royden Saah, Aaron B. Shiels, Paul Thomas, David W. Threadgill, Michael R. Vella, Fred Gould, Alun L. Lloyd

**Affiliations:** Department of Mathematics, North Carolina State University, Raleigh, NC 27695, USA; Biomathematics Graduate Program, North Carolina State University, Raleigh, NC 27695, USA; Department of Biological Sciences, North Carolina State University, Raleigh, NC 27695, USA; Island Conservation, 2100 Delaware Ave., Suite 1, Santa Cruz, CA 95060 USA; Department of Entomology and Plant Pathology, North Carolina State University, Raleigh, NC 27695, USA; National Wildlife Research Center, US Department of Agriculture, Fort Collins, CO 80521, USA; CSIRO Land & Water, Centre for Environment and Life Sciences, Floreat, WA, Australia; Genetic Engineering and Society Center, North Carolina State University, NC 27695, USA; Department of Microbiology, Immunology & Pathology, Colorado State University, Fort Collins, CO 80523, USA; School of Mathematical Sciences, The University of Adelaide, Adelaide, SA 5005, Australia; The Robinson Research Institute and School of Medicine, The University of Adelaide, Adelaide, SA 5005, Australia; Department of Molecular and Cellular Medicine, Texas A&M University, College Station, TX 77843, USA

## Abstract

Invasive species pose a major threat to biodiversity on islands. While successes have been achieved using traditional removal methods, such as toxicants aimed at rodents, these approaches have limitations and various off-target effects on island ecosystems. Gene drive technologies designed to eliminate a population provide an alternative approach, but the potential for drive-bearing individuals to escape from the target release area and impact populations elsewhere is a major concern. Here we propose the “Locally Fixed Alleles” approach as a novel means for localizing elimination by a drive to an island population that exhibits significant genetic isolation from neighboring populations. Our approach is based on the assumption that in small island populations of rodents, genetic drift will lead to multiple genomic alleles becoming fixed. In contrast, multiple alleles are likely to be maintained in larger populations on mainlands. Utilizing the high degree of genetic specificity achievable using homing drives, for example based on the CRISPR/Cas9 system, our approach aims at employing one or more locally fixed alleles as the target for a gene drive on a particular island. Using mathematical modeling, we explore the feasibility of this approach and the degree of localization that can be achieved. We show that across a wide range of parameter values, escape of the drive to a neighboring population in which the target allele is not fixed will at most lead to modest transient suppression of the non-target population. While the main focus of this paper is on elimination of a rodent pest from an island, we also discuss the utility of the locally fixed allele approach for the goals of population suppression or population replacement. Our analysis also provides a threshold condition for the ability of a gene drive to invade a partially resistant population.

## Introduction

Genetic modification of pest species has been suggested as a means to address a wide variety of pest problems, including those impacting human health, pre- and post-harvest crop losses, and conservation of endangered species^1–5^. One approach to genetic pest management involves introducing into the pest genome a DNA sequence that causes its own over-representation in future generations by inducing super-Mendelian inheritance; generally referred to as gene drive. An engineered gene coding for a desirable trait, e.g. one that renders the pest species less troublesome (e.g. reduction of vector competence of a mosquito species) can be linked to the gene drive sequence in order to increase its frequency in the population. Alternatively, the gene drive sequence can be engineered to disrupt a gene critical to fitness, or to deliver a payload that achieves this aim, and thereby suppress, or even eliminate, a population.

Ever since such drives were proposed, concerns have been raised in the peer-reviewed literature and in the popular media about unintended consequences of releases and ethical dimensions of the work. These include failure of the genetic construct being driven, but also the spread of gene drives beyond the region in which spread is intended and approved by the local population and governing authorities^2–13^. The nature and seriousness of these concerns differ between different gene drive technologies and applications (for instance, considerations would be quite different for an approach intended to replace a human disease-vectoring mosquito species by a variant that is refractory to the pathogen than for removal of an invasive species that causes ecological damage). These concerns are most acute in the case of drives that are designed to suppress and eliminate a population: spread of such a drive could have a risk of leading to global eradication of a species that is a pest in one area but a valuable component of an ecosystem in other areas (e.g. mice and rats). As a result, there is much interest in the ability to design gene drives that exhibit spatial localization, i.e. ones that have the ability to spread in a given region but will not spread globally.

Several approaches have been suggested to achieve localization of gene drives. Drives that exhibit an invasion threshold, such as the engineered underdominance (EU) approach, provide a natural means to achieve localization^14, 15^. These drives exhibit frequency-dependent dynamics where the drive can only spread if its frequency exceeds a particular level—the invasion threshold. Below this level, the frequency of the drive will decrease, leading to its loss from the population. Spread of a threshold drive across a patchy environment is more difficult, and becomes highly unlikely or even impossible when the invasion threshold is 50% or higher^16, 17^. Other approaches have been suggested to achieve localization, including Killer-Rescue^18^, Multi-locus assortment^19^, sex-linked genome editors^20^ and daisy-chain drive^21^. Theoretical analysis has suggested the ability of daisy-chain drive to simultaneously achieve spread and localization to a single area is only possible in a limited set of circumstances, and this concern also pertains to some other gene drives developed to be localized^22^. While all of these localized drives could change characteristics of pests in a population, their ability to locally suppress populations is questionable^22, 23^, although see also^24^.

In this paper, we propose a localization method, the “Locally Fixed Alleles” (LFA) approach, that can be utilized for relatively small populations that exhibit a significant degree of genetic isolation from other populations. This method is particularly suited for elimination of pest species from small oceanic islands, where the target population has small effective population size and for which there is naturally limited gene flow with other populations. While multiple alleles are expected to be commonly maintained at loci in large populations, genetic drift in small island populations is predicted to result in fixation of alleles at some loci in the genome^25, 26^.

Utilizing the high degree of genetic specificity of homing drives based on the CRISPR/Cas9 system^4^, our approach aims at employing one or more locally fixed alleles as the target for a gene drive on a particular island. Such a drive can spread to individuals carrying that allele, but individuals that do not have that specific allele are naturally resistant to the drive. For example, polymorphisms that occur in targeted Cas9 guide RNA binding sites or protospacer adjacent motifs (PAM) may effectively limit gene drive activity^27^ such that a drive can spread to individuals carrying alleles that form functional sites, but individuals with alternate alleles are naturally resistant to the drive. By design, we would search for alleles that are fixed in the population on the target island but not fixed in populations beyond that island. Consequently, the drive could be expected to result in only limited transient suppression beyond the island. A special case of this approach, the “private allele” (PA) approach (dubbed “precision drive” by Esvelt et al.^4^; see also^12^), occurs when the target allele is specific to the target population, but absent from other populations. We emphasize that the LFA approach does not require the target allele be a private allele, simply that it not be fixed in non-target populations.

The LFA approach can be used with a variety of different gene drives (e.g. standard homing drives, sex-biasing drives, and so on). Here, for simplicity, we illustrate the method using a standard homing drive^2, 4^ aimed at population elimination. We describe this in the setting of removal of a rodent species, such as the house mouse, *Mus musculus*, from an island. Invasive mice, and other rodents, are a particular concern for species conservation^28^, having significantly impacted many island ecosystems, including causing extinctions of endemic island vertebrate, invertebrate and plant species^29–31^. Although an island release would involve procedures that attempt to confine the gene drive mice to the island, unintended escape of these mice must be considered a possibility. Here, we use mathematical modeling to explore the impact that escape of drive individuals would have on mainland populations.

While the focus of this study is on a drive that can eliminate a rodent pest species from an island, the LFA approach can be used more generally for drives aimed at population suppression or replacement provided that the drive bears some fitness cost. These more general settings are discussed in detail in the Supplementary Information, including some important differences in the dynamics from the elimination setting discussed in the main text.

This paper is organized as follows: we first introduce the mathematical model, then briefly describe the single-patch (island-only) dynamics before discussing those seen in a two-patch (island-mainland) setting. We then explore the sensitivity of results to various drive and ecological/demographic parameters. Supplementary information includes the derivation of an analytic threshold condition for the ability of drive to invade a partially susceptible population, further sensitivity analyses, initial results of a stochastic model for the dynamics of LFA, and discussion of the use of LFA in more general population suppression and replacement settings.

## Methods

We employ a continuous-time non age-structured island-mainland model that describes the population dynamics and genetics of two populations. As our primary concern here is the impact of escape from the island, we assume unidirectional migration from the island to the mainland. (We recognize that migration from mainland to island would be an important consideration in the period following successful suppression or eradication from the island, but this is not the topic of this study.) Throughout, we assume a 50:50 sex ratio and so track numbers of female individuals. For an *n*-genotype system, denoting genotypes with a subscript and denoting island population numbers with superscript *I* (*N^I^_i_*) and mainland population numbers with a superscript *M* (*N^M^_i_*), we have

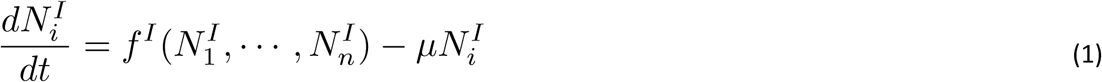

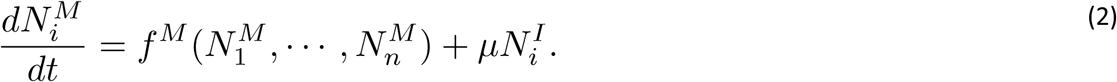

Here, the functions *f^I^* and *f^M^* describe the population dynamics and genetics that occur on island and mainland, respectively, and we assume unidirectional migration with per-capita migration rate equal to µ.

We assume that island and mainland populations both undergo random mating (i.e. are well-mixed) and exhibit logistic-type population dynamics. Our description of population dynamics is based on an earlier model^32^ for the population genetics and dynamics of gene drive in an island mouse population. Per-capita birth and death rates both change linearly with population size, with different coefficients on mainland and island (see^32^ and references therein). Within either the island or mainland, and in the absence of migration, genotype dynamics are described by

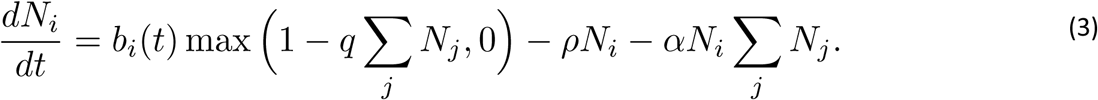

Here, superscripts denoting the location have been suppressed for clarity on both state variables and parameters. The functions *b_i_*(*t*), described below, depict the genotype-specific birth rates in the absence of density dependence. For an entirely wild-type population, *b*(*t*) would equal *λN*, where *λ* is the per-capita fecundity rate. The *b_i_*(*t*) are multiplied by a function that describes the linear density-dependent decline in per-capita birth rates with total population size. (Note that the max function is required to ensure that birth rates remain non-negative.) Per-capita death rates are assumed to increase linearly with overall population size but be independent of genotype. The density-independent component of the per-capita death rate (i.e. the reciprocal of the average lifespan when the population is at low density) is written as *ρ*, while the coefficient *α* describes the density-dependent linear increase in per-capita mortality. With these population dynamics, a single patch has a wild-type carrying capacity of *N* = *ρ*(*R*_0_ − 1)/(*λq* + α), where the basic reproductive number, *R*_0_, (i.e. the average number of female offspring of a female over its lifetime, at low population density) is equal to *λ*/*ρ*.

Population genetics is determined by the functions *b_i_*(*t*) which give the genotype-specific birth rates (c.f. the model of Robert et al.^33^) before accounting for the effects of density-dependence

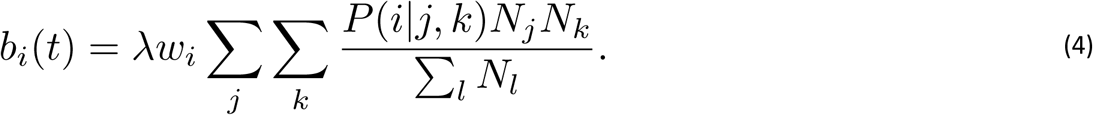

Here, *λ* is the baseline per-capita birth rate for females. P(*i*|*j*,*k*) gives the proportion of offspring from a mating involving individuals of types *j* and *k* that will have genotype *i*. The effects of the gene drive on biasing of inheritance are coded into these quantities. (The 216 entries of *P*(*i*|*j*,*k*), together with a fully written out set of model equations, appear in the Maple worksheet in the Supplementary Information.) The *w_i_* describe genotype-specific relative fitnesses, which we assume here to act at the embryonic stage and be equal in males and females of a given genotype.

As mentioned above, we use a simple homing-based elimination drive to illustrate the LFA method. We consider three alleles: the drive allele (D), the susceptible (S) allele, i.e. the target for the drive, and an allele that is resistant to the drive (R). All individuals in the target (island) population initially have the genotype SS, but those in the non-target (mainland) population can have SS, RS or RR genotypes. We assume only a single resistance allele in the mainland population. A large mainland population could have a number of different alleles that would be resistant to the drive, but they would all act similarly in being unaffected by the drive. Therefore our 3-allele model is sufficient for capturing dynamics of these cases. Note that here we are considering natural resistance to the drive, rather than drive-resistant alleles that are generated de novo as a result of the drive, e.g. by non-homologous end joining during homing. As a consequence, our model over-estimates the ability of the drive to spread and suppress populations^34^. Given that we are trying to evaluate the risk and implications of escape of drive, this assumption is conservative for our purposes. Homing is assumed to occur during gametogenesis, meaning that successful homing leads to an SD heterozygote individual giving rise to only D gametes (Homing exclusively in the germline at any point during development would have the same consequences). Successful homing occurs with probability *e*. The fitness, *w_i_*, of SS, RR, or SR individuals is assumed to be 1. The fitness of DD individuals is (1-*s*), and the fitness of SD and RD individuals is (1-*hs*), where *s* is the fitness cost of the drive and *h* is the degree of dominance of the fitness cost. For instance, a recessive lethal drive has *s* = 1 and *h* = 0. Notice that we assume there is no fitness cost for the RR individuals that occur naturally in the non-target population.

We employ parameters (see Table 1) that are appropriate for *Mus musculus* populations, largely based on those used by Backus & Gross^32^ (see also^35^). For this set of parameters, the basic reproductive number of a wild-type population is 3.5. Assuming an island of area 6 hectares, these parameters lead to an equilibrium population size of 1000 females. We assume that the mainland has a population size that is 100 times larger than this.

**Table 1:** List of Parameters and Their Values

|  | Parameter | Value | Source |
| --- | --- | --- | --- |
| $\lambda$ | Female fecundity rate | $8.4 \text{ (year)}^{-1}$ | Backus & Gross<sup>31</sup> |
| $q$ | Coefficient quantifying density-dependent decline in birth rate | Island: $6.427 \times 10^{-3} \text{ (mouse)}^{-1}$<br>Mainland: $6.427 \times 10^{-3} \text{ (mouse)}^{-1}$ | Backus & Gross<sup>31</sup> |
| $\rho$ | Density independent per-capita death rate | $2.4 \text{ (year)}^{-1}$ | Backus & Gross<sup>31</sup> |
| $\alpha$ | Coefficient quantifying density-dependent increase in death rate | Island: $5.000 \times 10^{-5} \text{ (mouse)}^{-1} \text{ (year)}^{-1}$<br>Mainland: $5.000 \times 10^{-7} \text{ (mouse)}^{-1} \text{ (year)}^{-1}$ | Backus & Gross<sup>31</sup> |
| $w_i$ | Genotype specific fitness | Dependent on $s$ and $h$ | |
| $\mu$ | Per-capita migration rate from island to mainland | Varied ($10^{-4}$-$1.2 \times 10^{-1} \text{ (year)}^{-1}$) | |
| $s$ | Drive allele fitness cost | 0.8 | |
| $h$ | Degree of dominance of fitness cost of drive allele | Non-threshold drive: 0.3<br>Threshold drive: 0.8 | |
| $e$ | Homing probability | 0.95 | |

## Results

### Single Patch Model

The single-patch behavior of the standard homing drive that we use here has been well-studied previously^2, 23, 36, 37^. Depending on the drive parameters *s, h* and *e*, several qualitatively different behaviors can occur following release of drive into an otherwise entirely susceptible population: guaranteed fixation of the drive, guaranteed loss of the drive, co-existence of drive and susceptible alleles at a stable polymorphic equilibrium or invasion threshold behavior (i.e. drive either goes to fixation or is lost, depending on its initial frequency) resulting from the existence of an unstable polymorphic equilibrium. Elimination of the population is possible when the drive remains in the population (going to a stable equilibrium with a positive frequency—either fixation or a polymorphic equilibrium) and imposes a cost that is sufficiently high to bring the reproductive number of the population below one.

We illustrate these dynamics using two sets of parameters: one for which the drive can spread through a susceptible population regardless of its frequency (no invasion threshold scenario), and another for which the drive can only spread when its frequency exceeds an invasion threshold (invasion threshold scenario). In both cases we consider a fitness cost of *s* = 0.8 and a homing probability of *e* = 0.95. The two scenarios differ only in the dominance, *h*, of the drive. For the no invasion threshold scenario we take *h* = 0.3, while for the invasion threshold scenario we take *h* = 0.8, which leads to an invasion threshold frequency of approximately 0.621 (corresponding to a ratio of approximately 1.64:1 drive:wild-type individuals). Releases occur into a population that is at carrying capacity, and population sizes are assessed relative to this carrying capacity.

Figure 1 shows the population dynamics that result from drive releases in these two scenarios. For the no invasion threshold scenario (green curve), even a small release of drive individuals, so that the initial relative population size is only just above one, leads to spread of the drive allele and hence reduction and eventual elimination of the population. For the invasion threshold scenario (blue and red curves), spread of the drive, and hence the fate of the population, depends on whether the release frequency of the drive is above (blue curve) or below (red curve) the invasion threshold. A sufficiently large release (blue curve) leads to fixation of the drive and elimination of the population, while an insufficient release (red curve) leads to loss of the drive and recovery of the population following a transient period of reduction.

**Figure 1:**
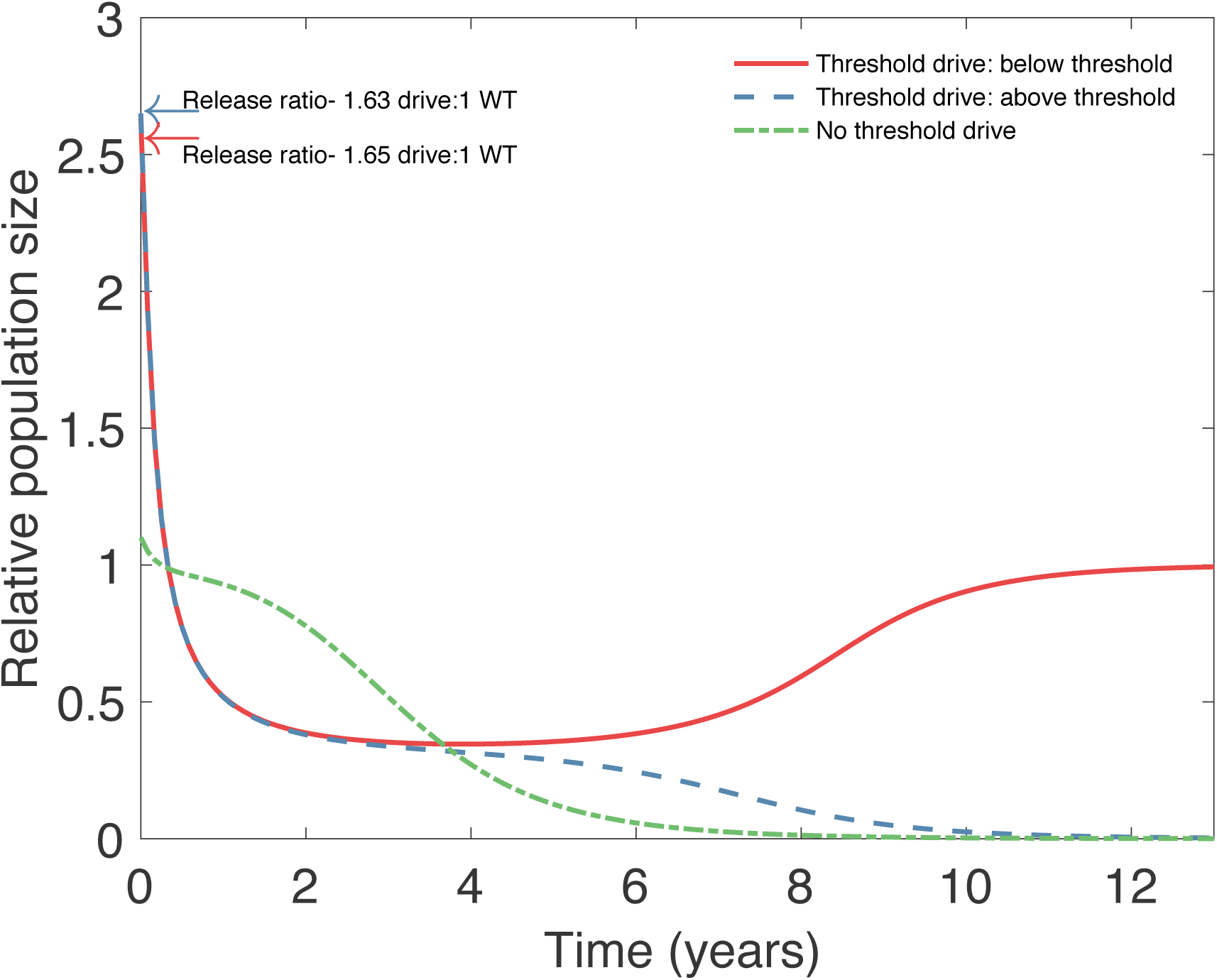
Island population dynamics for the no invasion threshold and invasion threshold scenarios. In both scenarios the drive has an 80% fitness cost (*s* = 0.8) but have differing degrees of dominance: for the no invasion threshold scenario (green curve) we take *h* = 0.3, while for the invasion threshold scenario we take *h* = 0.8. Population sizes are plotted relative to pre-release equilibrium population numbers. In the no invasion threshold scenario, arbitrarily small releases of drive individuals lead to invasion and fixation of the drive allele, leading to suppression of the population (green curve; release of 100 homozygous drive individuals, 0.1:1 release ratio). In the invasion threshold scenario, invasion of the drive depends on whether the initial release exceeds the invasion threshold (for this choice of parameters, the invasion threshold frequency for drive is 0.621, corresponding to approximately 1.64 drive individuals for each wild-type individual). Red curve depicts a sub-threshold release (1630 homozygous drive females; 1.63:1 release ratio), leading to loss of the drive allele and only temporary suppression of the population before its return to carrying capacity. Blue curve depicts a successful release (1650 homozygous drive females; 1.65:1 release ratio), for which the drive invades and reaches fixation, leading to elimination of the population. Values of other parameters are given in Table 1.

### Island-Mainland Model

We now turn to the main question of how migration from an island population on which drive individuals have been released will impact a mainland population that has a mix of susceptible and resistant individuals. To present something approaching a worst-case scenario, we first assume that the frequency of resistant individuals on the mainland is rather low, with a resistance allele frequency of 5% and susceptible allele frequency of 95%. We take our island population to be 1/100^th^ the size of the mainland population, with unidirectional migration from the island to the mainland occurring at per-capita rate of .012 per year, corresponding to movement at the rate of one mouse per month at baseline.

For the invasion threshold drive scenario, we consider an island release that is above threshold, so that the drive approaches fixation in the long run (Figure 2, upper panel, red curve) and the island population is successfully eliminated (Figure 2, upper panel, blue curve). Even though drive individuals migrate from the island to the mainland, the large size of the mainland population means that the drive frequency remains small and never exceeds the invasion threshold. Thus, the drive cannot spread on the mainland, leaving the size and genetic composition of the mainland population largely unaffected.

**Figure 2:**
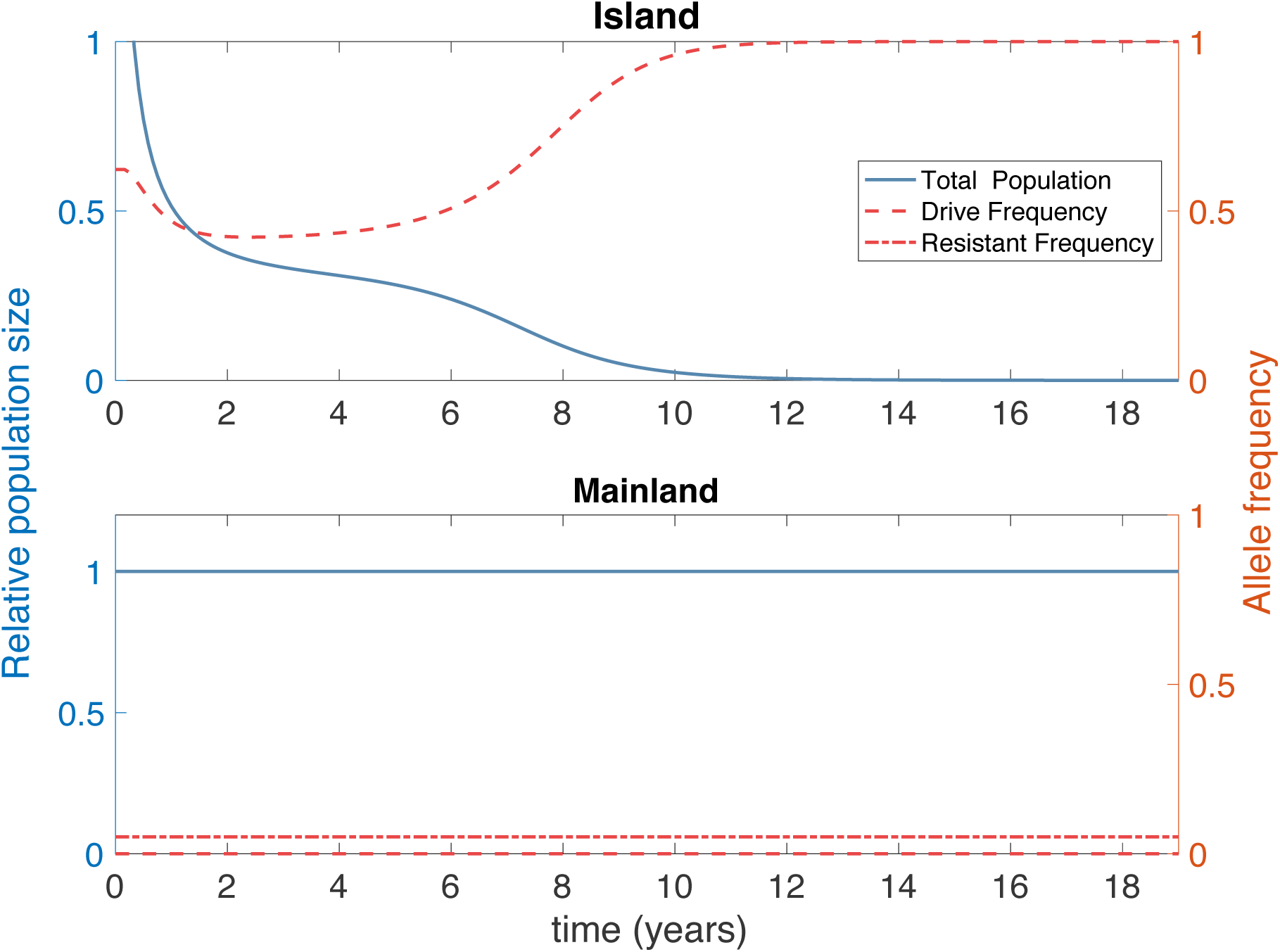
Island and mainland population dynamics (blue curves) and genetics (red curves) in the invasion threshold scenario (*s* = 0.8 and *h* = 0.8), following an above-threshold release of drive individuals on the island (1650 drive homozygotes released at *t* = 0; release ratio 1.65:1). Left axis (blue) denotes population size relative to pre-release equilibrium, right axis (red) denotes allele frequency. The drive spreads on the island (dashed red curve shows allele frequency for drive), suppressing its population (solid blue curve shows relative population size). Migration from the mainland to the island (on average, one island individual travels to the mainland a month) leads to a low drive frequency on the mainland that does not exceed the invasion threshold there. Consequently, the mainland population is largely unaffected. (Dot-dashed line denotes frequency of resistance allele. Note that the target allele being fixed means that the resistant allele is not present on the island.)

The situation is different in the no invasion threshold scenario. The drive is able to spread on the mainland even when present at the low frequencies that arise due to migration of drive individuals from the island. Individuals with a resistance allele are unaffected by the drive, however, and so spread only occurs through the susceptible individuals. Again, the drive spreads on the island (Figure 3, upper panel, red curve) causing successful elimination of the island population (Figure 3, upper panel, blue curve). On the mainland, drive spreads through the susceptible portion of the population (Figure 3, lower panel, red dashed curve) causing a reduction in the population size (Figure 3, lower panel, blue curve) due to fitness costs incurred by drive-bearing individuals. Because they are unaffected by the drive, resistant individuals benefit from having a larger relative fitness in the presence of drive individuals and so their frequency increases (Figure 3, lower panel, red dot dashed curve), while the drive frequency decreases. As the frequency of resistant individuals increases, the average fitness of the population returns to the level seen initially, and density-dependent dynamics returns the population to its original size. In this setting, we see a transient reduction in the mainland population size before a recovery due to the presence of resistance. The genetic composition of the mainland undergoes a shift during this process, with a reduction in the frequency of susceptible alleles (although not their elimination) and a corresponding increase in the frequency of resistance (see Supplemental Text and Supplemental Figure 3).

**Figure 3:**
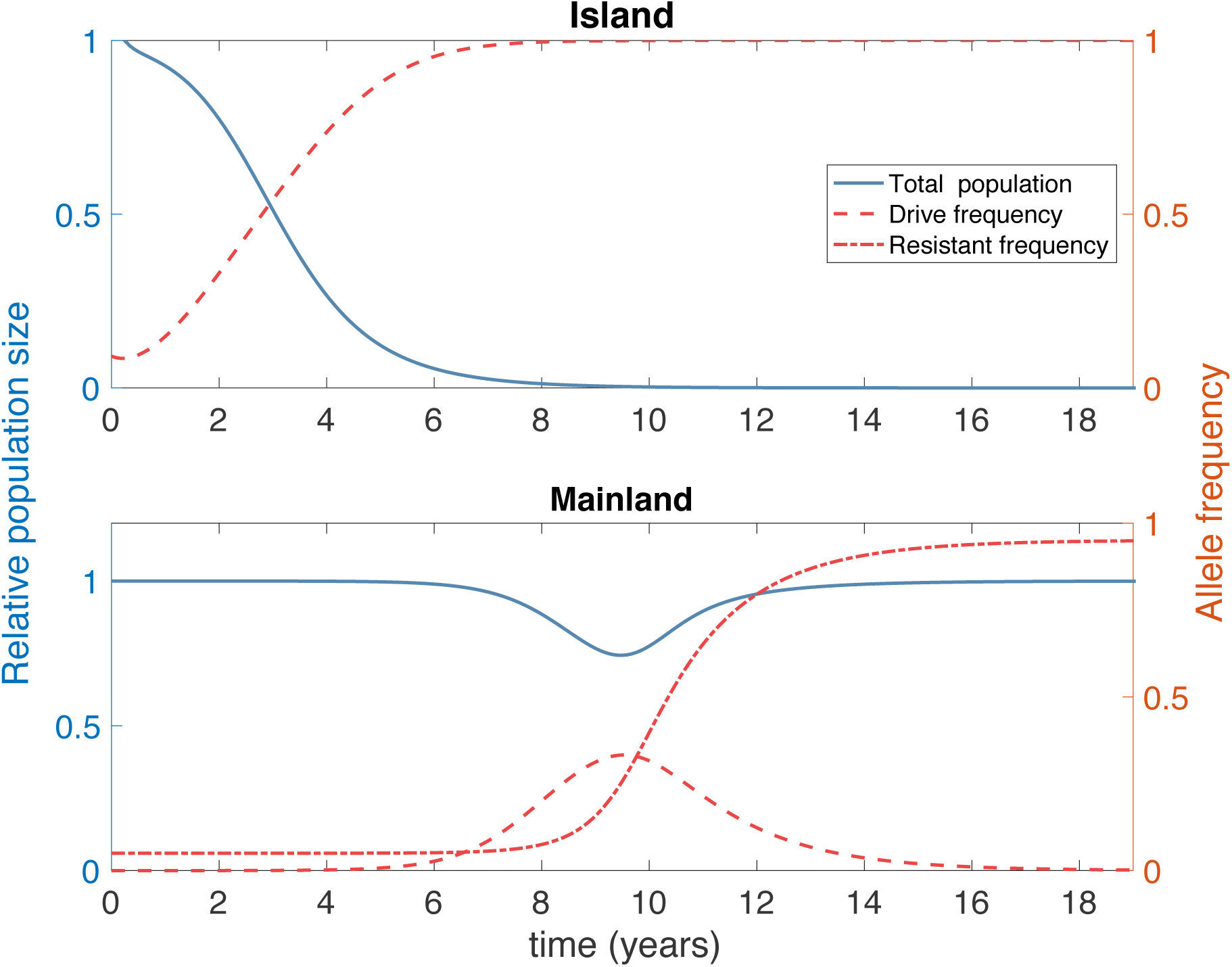
Island and mainland population dynamics and genetics in the no invasion threshold scenario (*s* = 0.8 and *h* = 0.3). As in Figure 2, blue curves and axes denote population sizes, measured relative to pre-release equilibria, while red curves and axes denote allele frequencies. One hundred homozygous drive individuals (0.1:1 release ratio) are released at time *t* = 0. We assume that resistance is very low on the mainland (allele frequency of just 5%), representing a rather pessimistic scenario in terms of the susceptibility of the mainland population to the drive. The drive spreads to fixation and suppresses the island population. Migration to the mainland (on average, one island individual travels to the mainland a month) means that the drive is introduced to the mainland, where it can spread through the susceptible population, but not the resistant population. The total population undergoes a temporary suppression as the drive spreads through the susceptible population. The frequency of resistant alleles increases as a result of drive, and density dependent population regulation returns the mainland population to the pre-release equilibrium level.

In the Supplementary Text we show that high levels of resistance prevent the spread of drive on the mainland. Using a linear invasion analysis, we show that drive cannot invade when the initial frequency of the resistant allele is above 1 - *hs*/{*e*(1-*hs*)}.

Figures 4 and 5 explore how the magnitude of population suppression and the maximum allele frequency of the gene drive observed on the mainland depend on the initial level of resistance on the mainland and the migration rate from the island to the mainland in the no invasion threshold scenario. Naturally, the level of transient suppression (fractional reduction below initial equilibrium) on the mainland depends strongly on the frequency of resistant and susceptible individuals there (Figure 4), although an important observation is that the level of transient suppression is considerably lower than the initial frequency of susceptible individuals. Similarly, the initial composition of the mainland population impacts the peak drive frequency reached on the mainland, while the level of migration again has little impact (Figure 5). For the assumed demographic and drive parameters, 10% or higher levels of resistance on the mainland lead to a transient population suppression of at most 20% and peak drive frequency below 30%. Alternative choices for demographic and drive parameters would impact these numbers (see sensitivity analyses in the Supplementary Information). Obviously, in the PA setting, drive would be unable to spread on the mainland, but Figures 4 and 5 reveal that even in situations for which the susceptible allele is not absent, but only fairly uncommon on the mainland, the levels of suppression that would result would be small, and quite likely smaller than natural fluctuations in population size that might result from demographic or environmental stochasticity.

**Figure 4:**
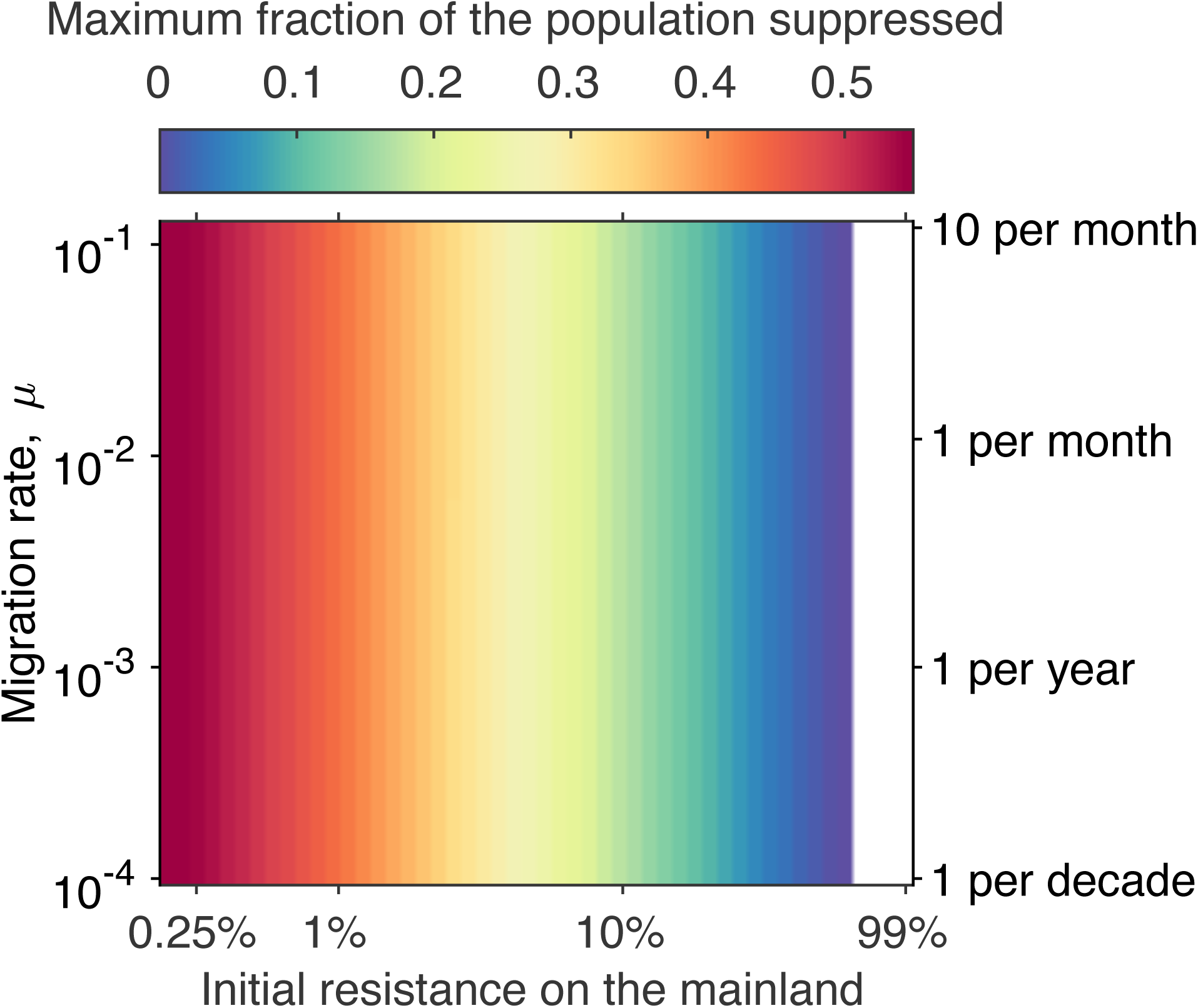
Maximum level of transient suppression seen on the mainland following an island release of 100 homozygous drive individuals in the no invasion threshold scenario (*s* = 0.8 and *h* = 0.3) across combinations of different migration rates and initial frequencies of resistant alleles on the mainland. Different levels of suppression are denoted by different colors (see color key on figure). Low initial resistance frequencies depict pessimistic scenarios (mainland is almost entirely susceptible to the drive), while high initial resistance frequencies approach the “private allele” scenario discussed in the text. The white region of the figure denotes initial levels of resistance that exceed the threshold level above which drive cannot invade the mainland.

**Figure 5:**
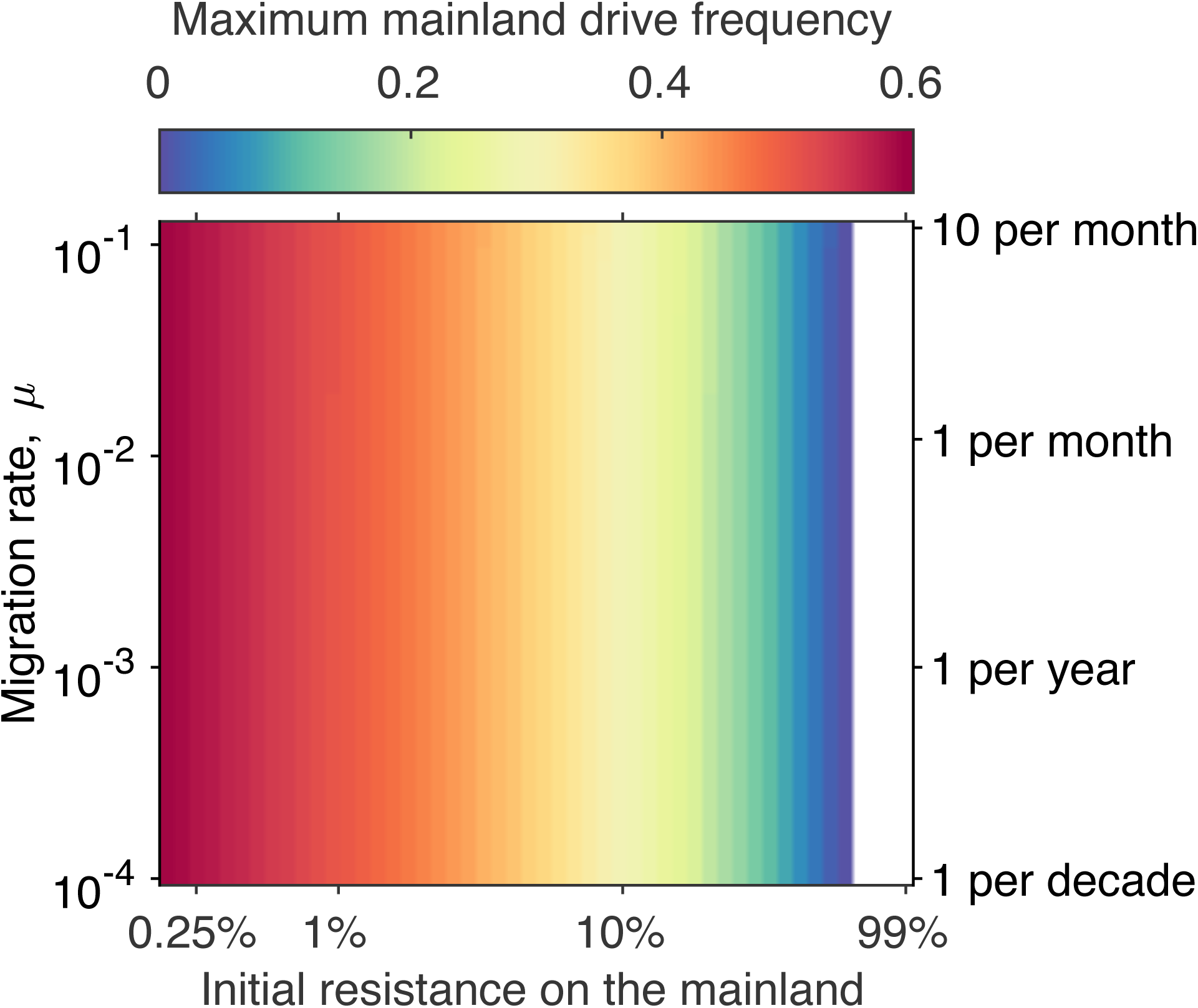
Maximum drive frequency observed on the mainland following an island release. Maximum drive frequency is indicated by color. All other details as in Figure 4.

The peak level of suppression and the maximum drive frequency on the mainland occur at roughly similar times. The timing of these events is relatively insensitive to the initial frequency of resistance on the mainland and the migration rate over a wide range of conditions (see Supplementary Text and Supplementary Figures 1 and 2), with dynamics playing out over the course of 10 to 20 years. High initial levels of resistance, however, can lead to much slower spread of the drive on the mainland. As mentioned above, drive cannot invade when the initial frequency of the resistant allele is above 1 - *hs*/{*e*(1-*hs*)}. Invasion of drive will occur only slowly (and with a low peak level) when the frequency of resistance is not too far below this level. Supplemental Figure 4 shows that the peak level of suppression on the mainland is almost independent of the size of the release on the island, while the time until this suppression occurs is only weakly dependent on the release size.

## Discussion

There is a fundamental tension between the ability of a gene drive to spread locally within a target area and its ability to invade populations beyond that area. For a non-threshold gene drive designed for suppression or elimination, the potential impact outside the targeted area can be of serious concern even if there is a likelihood that resistance to the drive will evolve before irreversible damage is done. Indeed, such concerns were strongly expressed in many of the seminal early papers on gene drive (e.g.^2, 3^). Clearly, mechanisms that give some level of control over the spread of a gene drive are prerequisites for the deployment of a gene drive for suppression or elimination of a targeted population if other populations of the species contribute positively to biodiversity or economics. Utilizing the high degree of genetic specificity exhibited by homing drives, the LFA approach provides one option to limit the impact of unintended spread of a drive. In the fortunate, although perhaps rather unlikely special case of “private alleles” where all individuals in non-target populations lack the susceptible allele and the susceptible allele is fixed on the island, the drive would be completely confined to the target population.

The LFA approach is related to two-step gene drive approaches that first spread a target allele into a population and then release a second drive that is specific to the target allele. For example, Esvelt *et al.*^4^ suggested this approach for island populations, arguing that containment would happen provided that release of the second drive occurred before any individuals bearing the first drive escaped the island. The uncertainty involved in this scenario could be problematic for some stakeholders and regulatory authorities. In our approach, naturally occurring targeted alleles are expected to be fixed within small island populations due to genetic drift or founder effects that have occurred in the past. These targeted alleles may be present in other locations, but as long as they are not fixed, impacts on populations in those locations should be transient.

As demonstrated by other mathematical models (e.g.^22, 38^) some spatially limited gene drive mechanisms are unlikely to function well for strongly suppressing populations, because these require extremely large releases, which are neither feasible nor advisable for pest species on islands (but see^39^ for an alternative approach for localized suppression on mainland populations). In this regard, the LFA approach stands out for its predicted ability to efficiently eliminate a small mostly isolated population without impacting other populations beyond a transient effect. The LFA approach could also be useful in the setting of a suppression or replacement drive that does not lead to elimination of the island population, although, as discussed in detail in the Supplementary Information (Section S.5), persistence of the drive on the island will mean continual reintroductions of drive to the mainland via migration, most likely leading to maintenance of drive on the mainland at a low frequency.

Our analysis employed a deterministic model, which allows for continuous migration from island to mainland. This model describes the average behavior of the system and so gives a good indication of expected dynamics in a large well-mixed mainland population. It describes the magnitude of suppression and its timing, but does not account for the discrete numbers of individuals in populations. Given that migration from the island to mainland is anticipated to occur as relatively infrequent events that involve small numbers of individuals, a stochastic model would be more realistic. Such a model would allow questions about the likelihood of the occurrence of migration events and of resulting transient spread of the drive, and distributions of the magnitude and timing of any transient suppression that results. However, we would not expect the qualitative results of such a model to predict a qualitatively different outcome from the current model in terms of impacts on the overall mainland population (see Supplementary Information for results from a stochastic model and some additional discussion of the impact of stochasticity).

For this proof-of-concept study of the LFA approach, we employed a highly simplified description of the population dynamics and genetics of the system. Many ecological and behavioral complexities, such as Allee effects^40^, the spatial and social structures of mouse populations, and mouse mating behavior, will impact the dynamics of population suppression and elimination under the action of gene drives. More refined models that include such features will have to be developed if the use of this approach is to be considered for a real-world release program.

It is important to recognize that one of the outcomes of mice arriving on the mainland that have a homing drive with no threshold is that the frequency of the resistant allele will increase. If after the mice are eliminated from the island a mouse or a few mice carrying the now more common resistant allele migrate to the island, it is likely that the new population on the island would no longer be a good target for the previously used construct. However, if recolonization was started by one or a few individuals, founder effects and drift would be likely to result in other fixed alleles that could be targeted.

There are a number of technical and societal challenges in moving the LFA from concept to application. While there is empirical evidence of lower polymorphism in small island populations (e.g.^41–43^), not every fixed allele in an island population will be a good target for a CRISPR-based gene drive. Appropriate targets will depend on the intended nature of the drive (e.g. if it is to be sex-specific, employ a split-drive design, etc), impacting the number of available loci. Furthermore, these alleles will optimally have at least two sites that can be targeted by the CRISPR-CAS nuclease complex to decrease the chance of drive failure due to resistant alleles arising through non-homologous end joining^44^. It will be critical that full genomes of individuals be scanned for the best possible targets and those targets will need to be scrutinized for potential problems. Also, because the LFA approach requires that the target allele is fixed, island populations will need to be sampled and evaluated extensively before moving ahead with any genetic engineering to ensure an acceptably high probability that the targeted allele is truly fixed on the island. This will require focused sequencing of the target loci in a large number of individuals, but it should be recognized that 100% assurance is not possible.

At a societal level, lack of an ecological impact of an LFA in a non-target location of the specific target species must be assured. Models that include details of the species biology, population structure and genetics will be critical in moving toward regulatory approval. But assurance of lack of an ecological impact of LFA constructs on the non-targeted populations may not be sufficient for some stakeholders who will have a concern with even one engineered individual arriving in their area.

Beyond direct impacts of an LFA, researchers must be cognizant of the fact that in developing the technology for LFA in a problematic species, they are also developing tools that could be used by others to construct unrestricted gene drives in that species. There will always be tradeoffs in developing new technologies and there are unlikely to be simple decisions. Vigilance and input from diverse stakeholders “early and often”^45^ will be critical in coming to decisions.

## Acknowledgements

This work was funded by the DARPA Safe Genes Program (Grant DARPA-16-59-SAFE-FP-005), from which several of the authors (DVB, KJC, SD, JDE, OE, JG, GRH, KO, AJP, TAAP, ABS, PT and ALL) receive support. Support was also received from the Research Training Group in Mathematical Biology, funded by NSF grant RTG/DMS-1246991 (JS, BH, MRV and ALL), NSF IGERT grant 1068676 (MRV, FG and ALL), NIH grant 1R01AI139085-01 (FG and ALL) and the NC State Drexel Endowment (ALL). Contributions from other members of the Genetic Biocontrol of Invasive Rodents (GBIRd) consortium (http://www.geneticbiocontrol.org/) are acknowledged and greatly appreciated. We thank the referees for their constructive comments that helped improve this paper.

## Author Contributions

JS, BH, FG and ALL designed the model. JS, BH and ALL carried out model simulations and analysis. JS, BH, FG and ALL wrote the first draft of the paper. All authors discussed model results and contributed to editing and revision of the manuscript.

## Supplementary Information

### S.1. Threshold for Invasion of Drive into Partially Susceptible/Partially Resistant Mainland Population

It can be shown that there is a threshold level of resistance above which drive cannot invade the mainland. This threshold can be derived by examining the stability of a drive-free equilibrium state of the single patch model, using the standard approach of linearizing the model about the equilibrium (see, for example^1^). This process involves calculation of the eigenvalues of the Jacobian matrix (i.e. the matrix of the partial derivatives of the right-hand sides of the differential equations (eqns 3 in the main text) with respect to the state variables of the model) evaluated at the equilibrium of interest. The 6×6 Jacobian matrix and its eigenvalues are easily calculated with the assistance of a computer algebra package (e.g. Maple [Maplesoft, a division of Waterloo Maple, Inc., Waterloo, Ontario] or Mathematica [Wolfram Research, Inc., Champaign, IL]). A Maple worksheet providing details of the calculations is provided as a supplementary file.

Given that we assume equal fitness of susceptible and resistant types, in the absence of drive there is no selection between susceptible and resistant types and so we have non-unique drive-free equilibria:

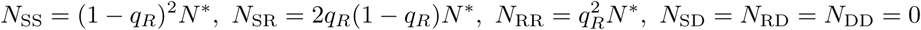

Here *q*_R_ is the frequency of the resistant allele and *N*\* is the wild-type (i.e. drive-free) equilibrium population size (equal to (*λ*-*ρ*)/(*λq*+*α*)). We note that these equilibria form a curve in phase space.

The linear stability analysis gives six eigenvalues, one of which is zero (corresponding to neutral stability along the curve of drive-free equilibria), one equal to *ρ*-*λ* and three equal to –*λ*(*qρ*+*α*)/(*λq*+*α*). We see that these last four eigenvalues are negative (note that *λ* must be greater than *ρ* in order for *N*\* to be positive). The ability of drive to invade (from low initial frequency) is determined by the sign of the final eigenvalue, whose value is equal to *λ*[(1-*q*_R_)(1-*hs*)*e*-*hs*](*qρ*+*α*)/(*λq*+*α*). Invasion is only possible if this quantity is positive, and so consideration of the quantity in square brackets leads to the condition

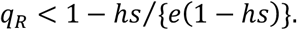

Another way to derive this threshold (and one that immediately applies to a number of previous studies in the literature) is by considering a discrete-time (discrete generations) description framed in terms of allele frequencies (see, for example^2, 3^).

If the allele frequencies for Susceptible, Drive and Resistant alleles in the current generation are written as *q*_S_, *q*_D_ and *q*_R_, it can be shown that the frequency of the drive allele in the next generation, *q*′_D_, will be given by

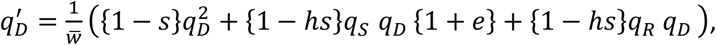

where 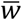 is the mean fitness, which equals

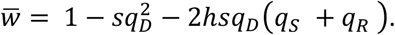

When thinking about invasion of drive into a population that initially consists of susceptible and resistant alleles, *q*_D_ will be small, and so the following linear equation can be derived for the change in the frequency of the drive allele from one generation to the next:

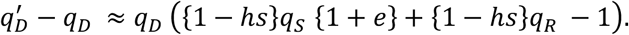

(This equation is correct to first order in *q*_D_.)

We see that the drive frequency can only increase if the term in parentheses is positive, meaning that

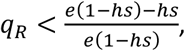

which may be written as

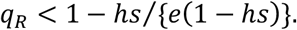

### S.2. Additional Details of Dynamics on the Mainland: Timing of Suppression and Peak Drive Level; Long-Term Dynamics

Supplemental Figures 1 and 2 explore the timing of (1) suppression on the mainland and (2) the peak level of drive seen on the mainland. These two events occur at similar times (but not at exactly the same time). We note that over a wide range of initial levels of resistance and migration rates, there is only weak dependence of either time on these quantities, and that these times increase substantially as the initial level of resistance approaches the non-invasion threshold.

Supplemental Figure 3 explores the composition of the mainland population that remains after the transient spread of drive, showing the long-term (100 year) frequency of the susceptible allele. We note that, while the transient spread of drive leads to the reduction of susceptible alleles on the mainland, it does not lead to their elimination.

Supplemental Figure 4 shows that the peak level of suppression and the maximum drive frequency on the mainland are almost independent of the size of the release on the island (that they are not constant cannot be seen at the scale shown on this figure: both curves exhibit a very weak dependence on release size). The time until the peak suppression occurs is only weakly dependent on the release size for biologically plausible release sizes, varying only by about 3 years for release sizes between 1 and 1000 individuals, but continuing to increase in the limit as the release size approaches zero. In the deterministic model, invasion is possible from arbitrarily small releases, although it takes increasingly long for drive to increase to appreciable levels from extremely small release levels, hence the time to minimum continues to increase. Note that the model behaves qualitatively differently when the release size is zero from when the release size is non-zero.

**Supplemental Figure 1.**
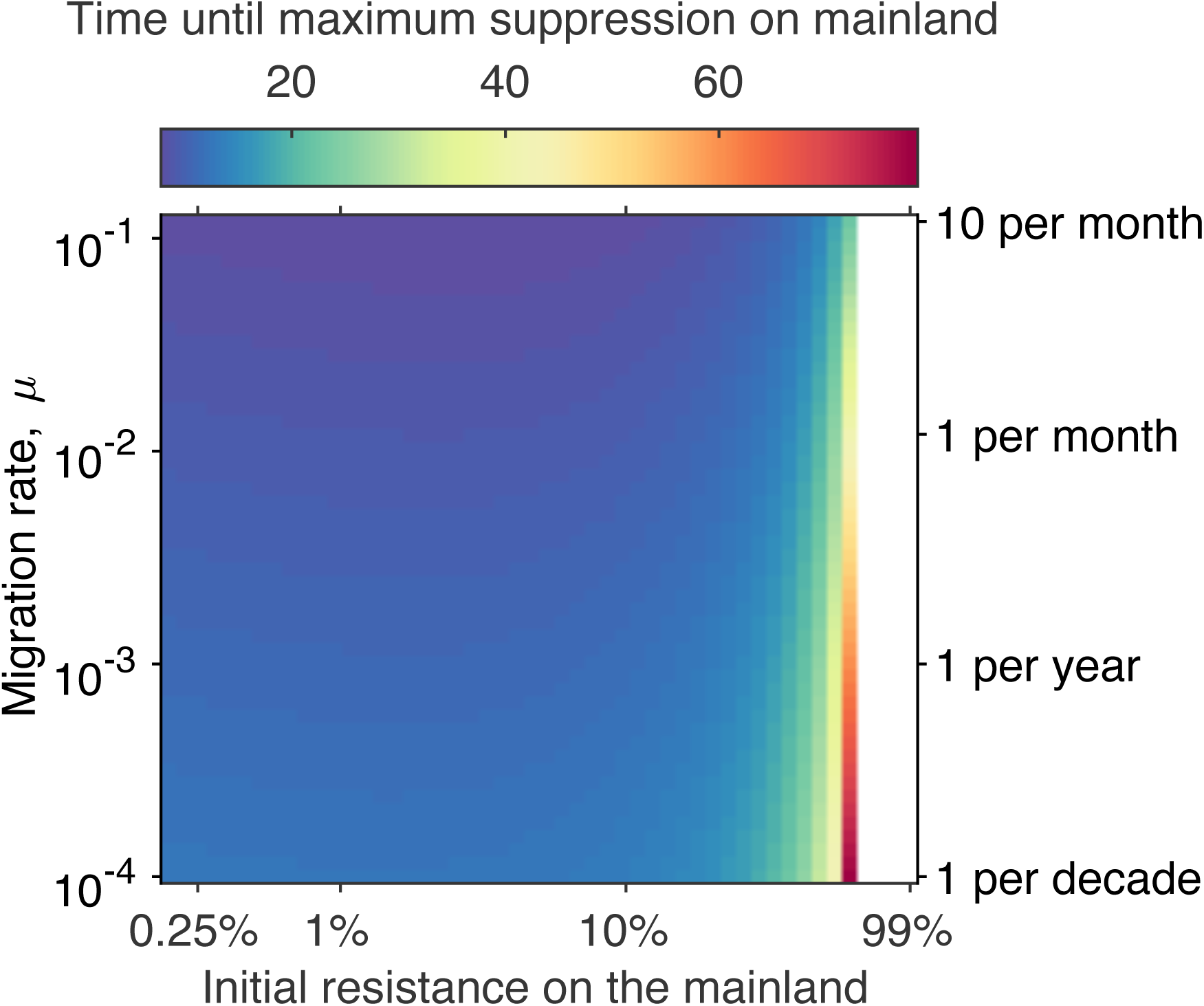
Dependence of the time until the mainland population achieves its minimum on the initial level of resistance on the mainland and the migration rate from the island. All other details are as in Figure 4 of the main text. The white region of the figure denotes initial levels of resistance that exceed the threshold level above which drive cannot invade the mainland.

**Supplemental Figure 2.**
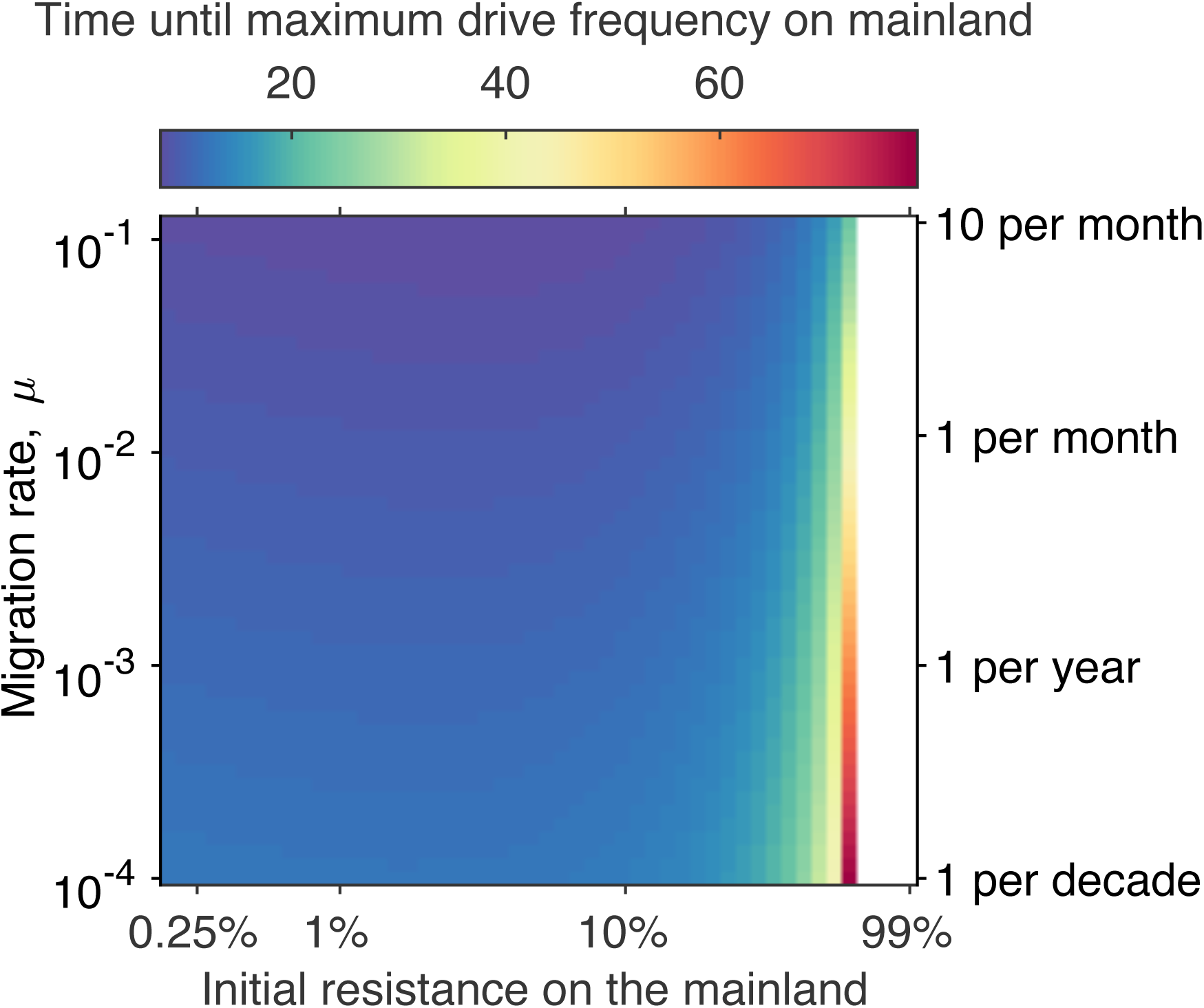
Dependence of the time until the drive frequency achieves its maximum on the mainland on the initial level of resistance on the mainland and the migration rate from the island. All other details are as in Figure 4 of the main text.

**Supplemental Figure 3.**
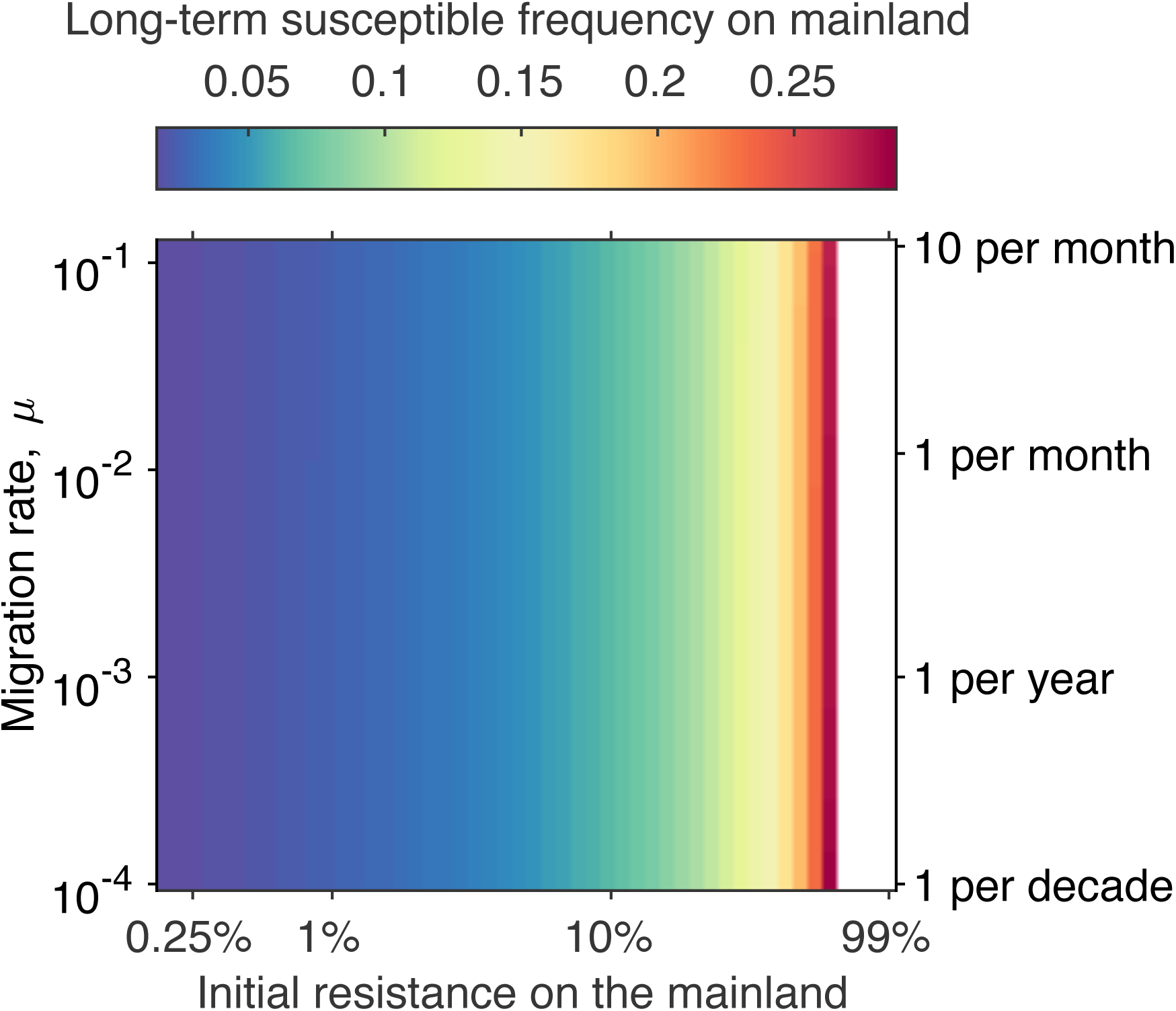
Frequency of the susceptible allele on the mainland 100 years after an island release in the no invasion threshold scenario (*s* = 0.8 and *h* = 0.3) across combinations of different migration rates and initial frequencies of resistant alleles on the mainland. All other details are as in Figure 4 of the main text. Note that susceptible alleles remain in the population after the transient spread and loss of drive on the mainland. The white region of the figure denotes initial levels of resistance that exceed the threshold level above which drive cannot invade the mainland.

**Supplemental Figure 4.**
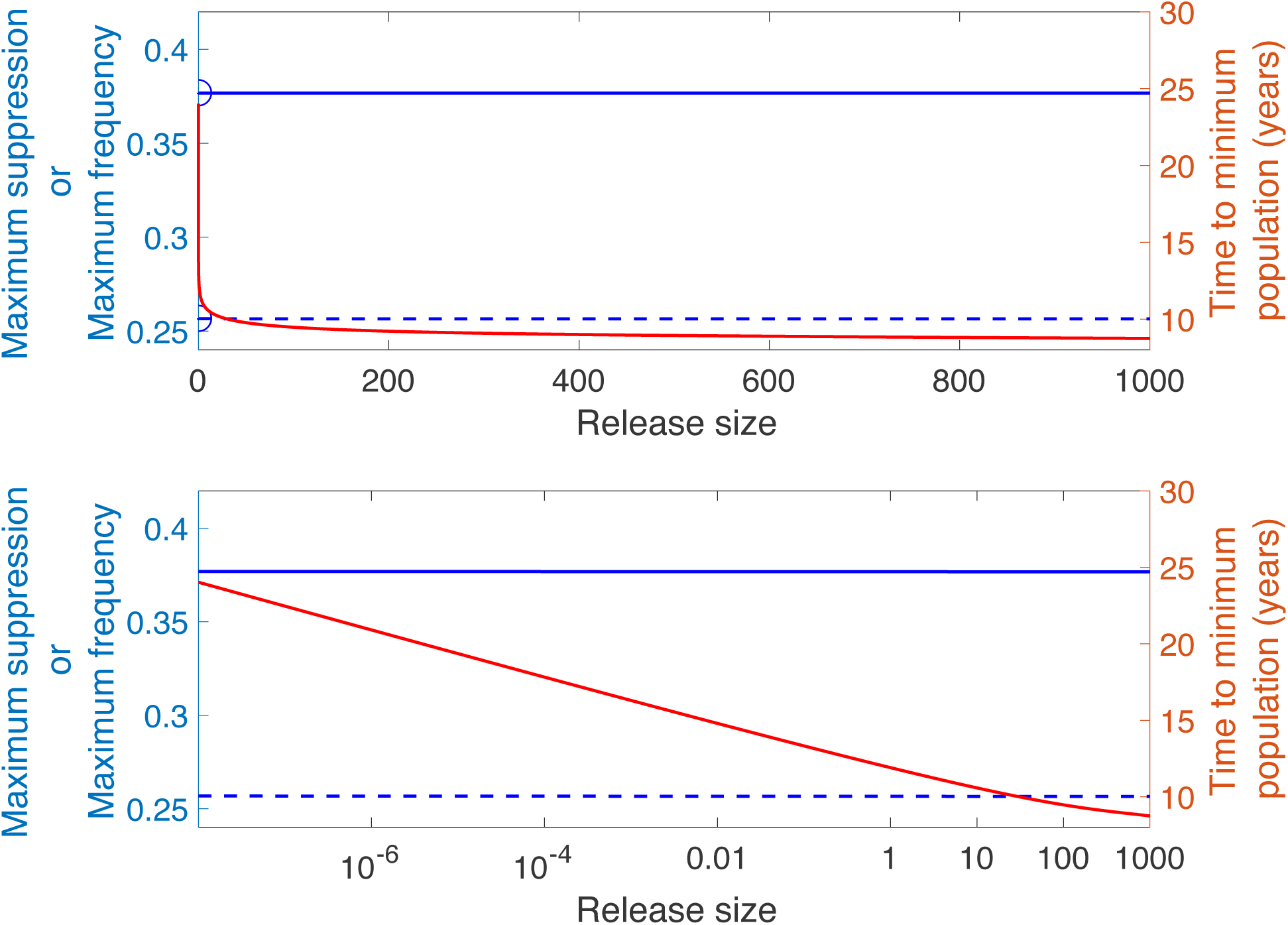
(Both panels) Maximum level of transient suppression (dashed blue curve; read scale on left axis), time taken for this level of suppression to occur (red curve, read scale on right axis) and maximum level of gene drive (solid blue curve; read scale on left axis) seen on mainland following releases of various sizes on the island under the no invasion threshold scenario. Initial frequency of resistance on the mainland is equal to 0.05 and migration occurs at a per-capita rate of 0.012 per year. All other parameters are as in Figure 3 of the main text. Circles on the vertical axis denote that the curves are not continuous when the release size is zero: the behavior of the model is qualitatively different between zero and positive release sizes. The bottom panel depicts the same results but using a logarithmic scale on the horizontal axis to emphasize the mathematical behavior as the release size approaches zero.

### S.3. Sensitivity of Results to Parameter Values

In this section we explore how model results, specifically the maximum level of suppression seen on the mainland and the peak level reached by drive on the mainland, depend on the drive and ecological parameters used. Such a sensitivity analysis provides increased confidence in the utility of the LFA approach.

Before exploring the impact of individual parameters in detail, we performed a global sensitivity analysis involving four parameters: the fitness cost of the drive, *s*, the dominance of this fitness cost, *h*, the homing probability, *e*, and the demographic parameter *λ*, the female fecundity parameter (i.e. the female per-capita birth rate of the population at low population densities, when density-dependent reductions in birth rate can be ignored). As discussed below (S.3.2.1), for the logistic model, if we choose to fix the equilibrium population size, there is only one demographic parameter, the net per-capita birth/death rate at low population densities, that can be independently varied. We choose to do this by varying the per-capita birth rate (female fecundity, *λ*).

Given that we have little to no information on the uncertainties of these parameters about their baselines, we employed uniform distributions for their possible values (Table S.1), following the approach taken by Prowse et al.^4^ in their sensitivity analysis of the Y-CHOPE Y chromosome shredding gene drive. We then used a sampling-based variation decomposition approach^5^ (Fourier Analysis Sensitivity Test, FAST, implemented using the SAFE Toolbox^6^) to assess the contributions of the uncertainties in different parameters to the variation seen in model outputs across the parameter space. This approach is somewhat akin to more familiar analysis of variance statistical methods. 5000 simulation runs were carried out, each using a set of parameters sampled from the parameter space described in Table S.1.

**Supplemental Table 1:** Baseline values and assumed distributions for parametric sensitivity analysis.

| Parameter | Baseline | Distribution |
| --- | --- | --- |
| $s$ | 0.8 | U(0.65,0.95) |
| $h$ | 0.3 | U(0.1,0.5) |
| $e$ | 0.95 | U(0.7,1.0) |
| $\lambda$ | 8.4 | U(6,10) |

Across the simulation runs based on 5000 sets of parameters, the range of observed maximum suppression values fall between less than 1% and 40.4%, with a mean of 20.8% and standard deviation of 0.08% (coefficient of variation of 38.8%). For the maximum drive frequency, the range of observed values was between less than 1% and 53.7%, with a mean of 31.8% and standard deviation of 12.2% (coefficient of variation of 38.2%).

For maximum suppression observed on the mainland, the first order effects of the four parameters explained 84% of the variation. Two of the drive parameters, *h* and *e*, and the demographic parameter *λ* explained fairly similar amounts of variation (26%, 22%, and 23%, respectively), while the fitness cost *s* explained just 12% of the variation. This analysis says that if we wished to predict the impact of LFA on mainland population suppression, reducing the uncertainty in either *h*, *e*, or *λ* has a bigger impact on the confidence in our predictions than reducing uncertainty in *s*.

For maximum drive frequency observed on the mainland, the first order effects of the four parameters explained 94% of the variation. The drive fitness parameters *s* and *h* have the biggest impact, explaining 47% and 38% of variation, respectively. The homing probability *h* has a much smaller impact, explaining 8% of variation, while the demographic parameter *λ* has very little impact on the maximum drive frequency, explaining less than 0.1% of variation (see S.3.2.1 for more discussion on this).

#### S.3.1. Sensitivity to Drive Parameters

Figure S.5 shows the dependence of the maximum levels of suppression and drive seen on the mainland on the drive fitness cost and dominance of this fitness cost. Results are shown for the region of drive parameter space for which the island population is eliminated and for which no threshold behavior is observed (Deredec et al.^2^ and Alphey and Bonsall^7^ provide analytic expressions for the locations of these boundaries). The dashed line on the plot shows the boundary between the parameter regions for which drive becomes fixed or approaches a polymorphic equilibrium with wild-type^2, 7^.

Figure S.6 shows sensitivity of outcomes over a region of drive parameter space assuming different values for the homing probability, and Figure S.7 shows dependence of outcomes on the initial level of resistance on the mainland assuming different values for the homing probability.

**Supplemental Figure 5:**
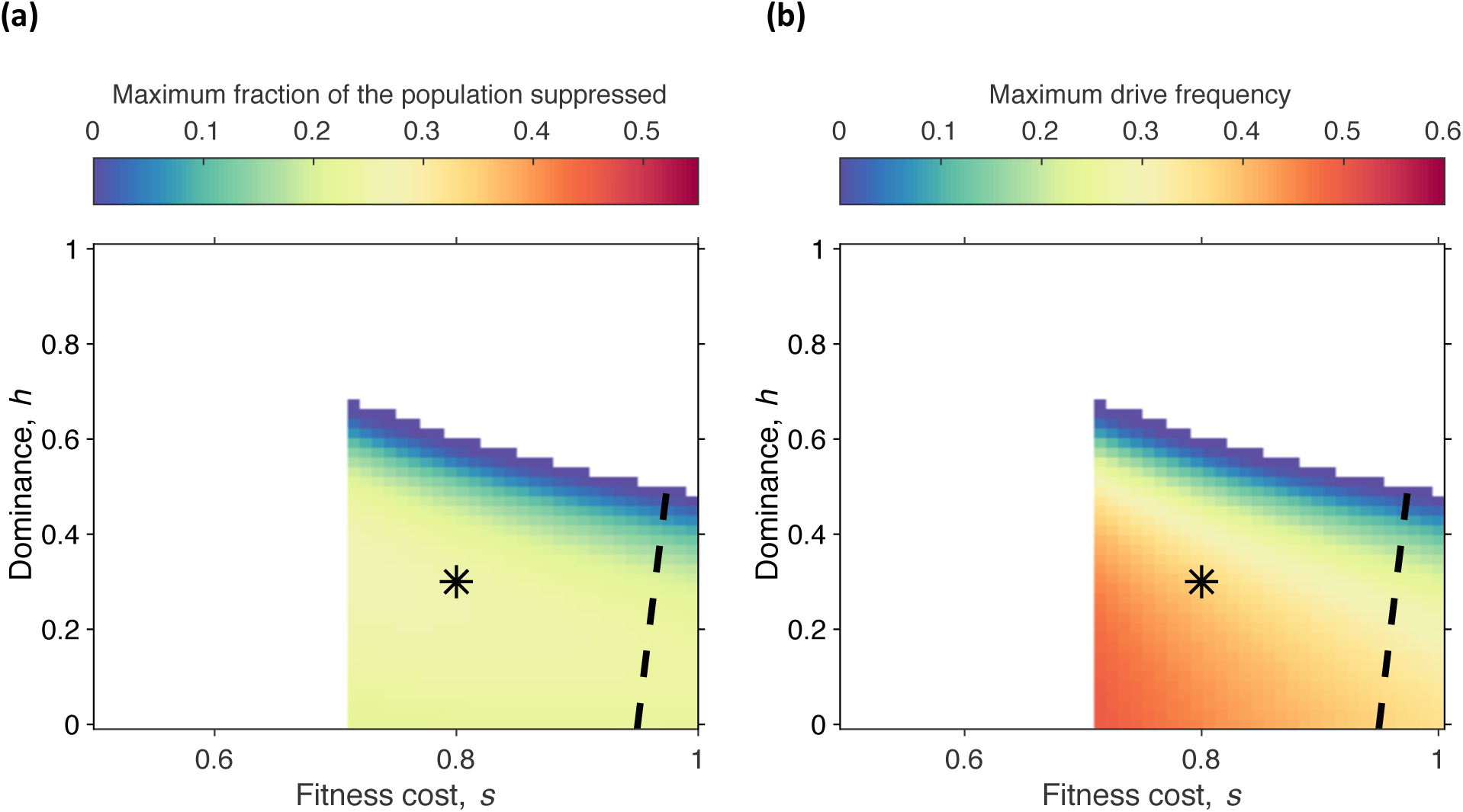
Heatmaps showing the dependence of (panel a) the maximum suppression observed on the mainland and (panel b) the maximum level of drive seen on the mainland on the drive parameters *s* and *h*. The scales on the color bars are chosen to be the same as in Figures 4 and 5. White regions on this figure denote combinations of drive parameters that either lead to threshold behavior, loss of drive or for which the drive fails to lead to elimination of the island population. The initial frequency of the resistance allele on the mainland is 95%, and the migration rate is 0.012/year. All other parameters are as in Figures 4 and 5 of the main text. The black asterisk denotes the drive parameters used in Figures 4 and 5 of the main text. The dashed line denotes the boundary between the region of parameter space where drive approaches fixation (to the left of the line) and where drive approaches a polymorphic equilibrium with wild-type (to the right of the line).

**Supplemental Figure 6:**
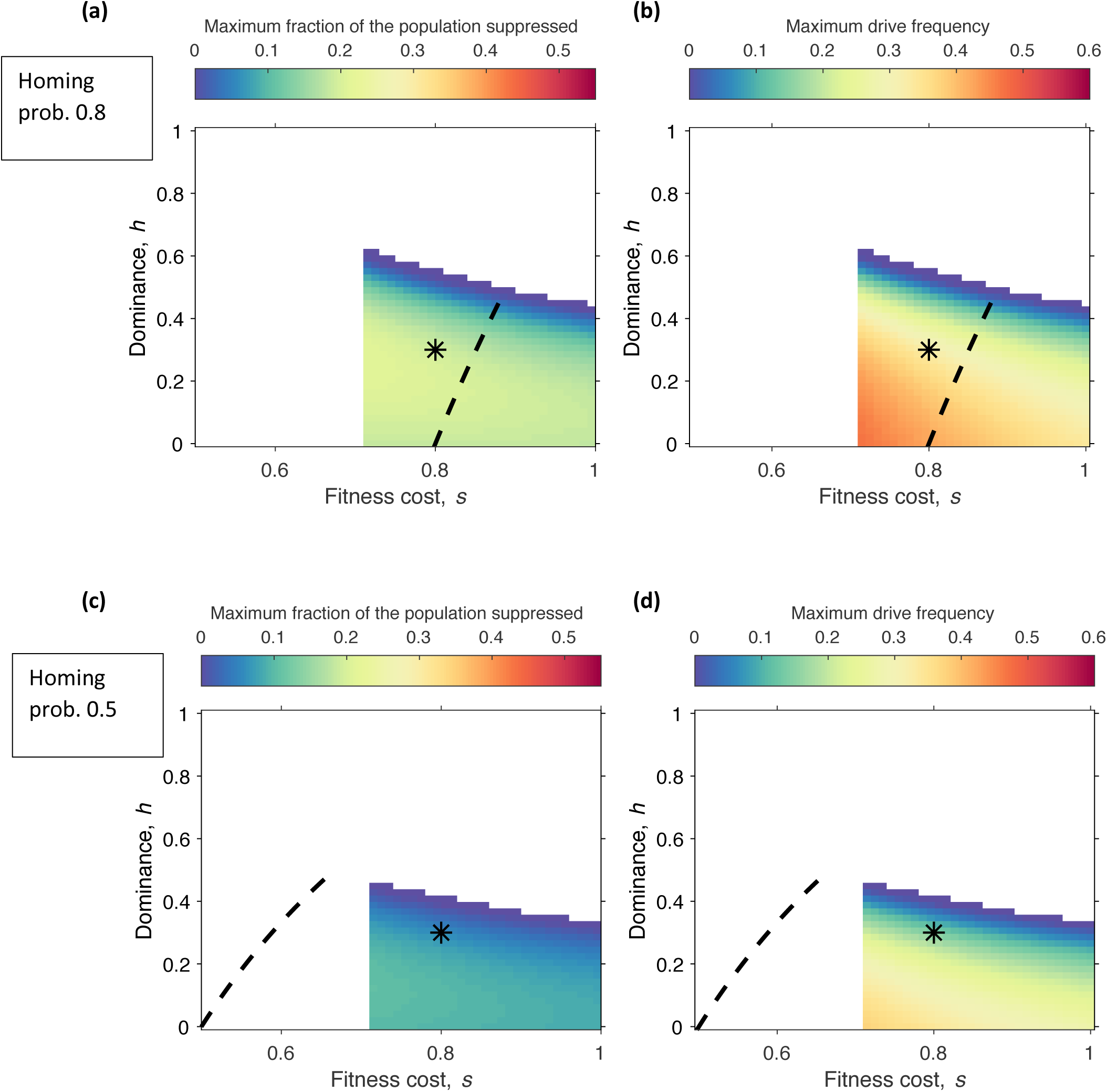
Heatmaps showing the dependence of (panels a and c) the maximum suppression observed on the mainland and (panels b and d) the maximum level of drive seen on the mainland on the drive parameters *s* and *h*, and for homing probabilities of (panels a and b) 0.8 and (panels c and d) 0.5. The scales on the color bars are chosen to be the same as in Figures 4 and 5 of the main text. White regions on this figure denote combinations of drive parameters that either lead to threshold behavior, loss of drive or for which the drive fails to lead to elimination of the island population. The initial frequency of the resistance allele on the mainland is 95%, and the migration rate is 0.012/year. All other parameters are as in Figures 4 and 5 of the main text. The black asterisk denotes the drive parameters used in Figures 4 and 5 of the main text. The dashed curve denotes the boundary between the region of parameter space where drive approaches fixation (to the left of the curve) and where drive approaches a polymorphic equilibrium with wild-type (to the right of the curve).

**Supplemental Figure 7.**
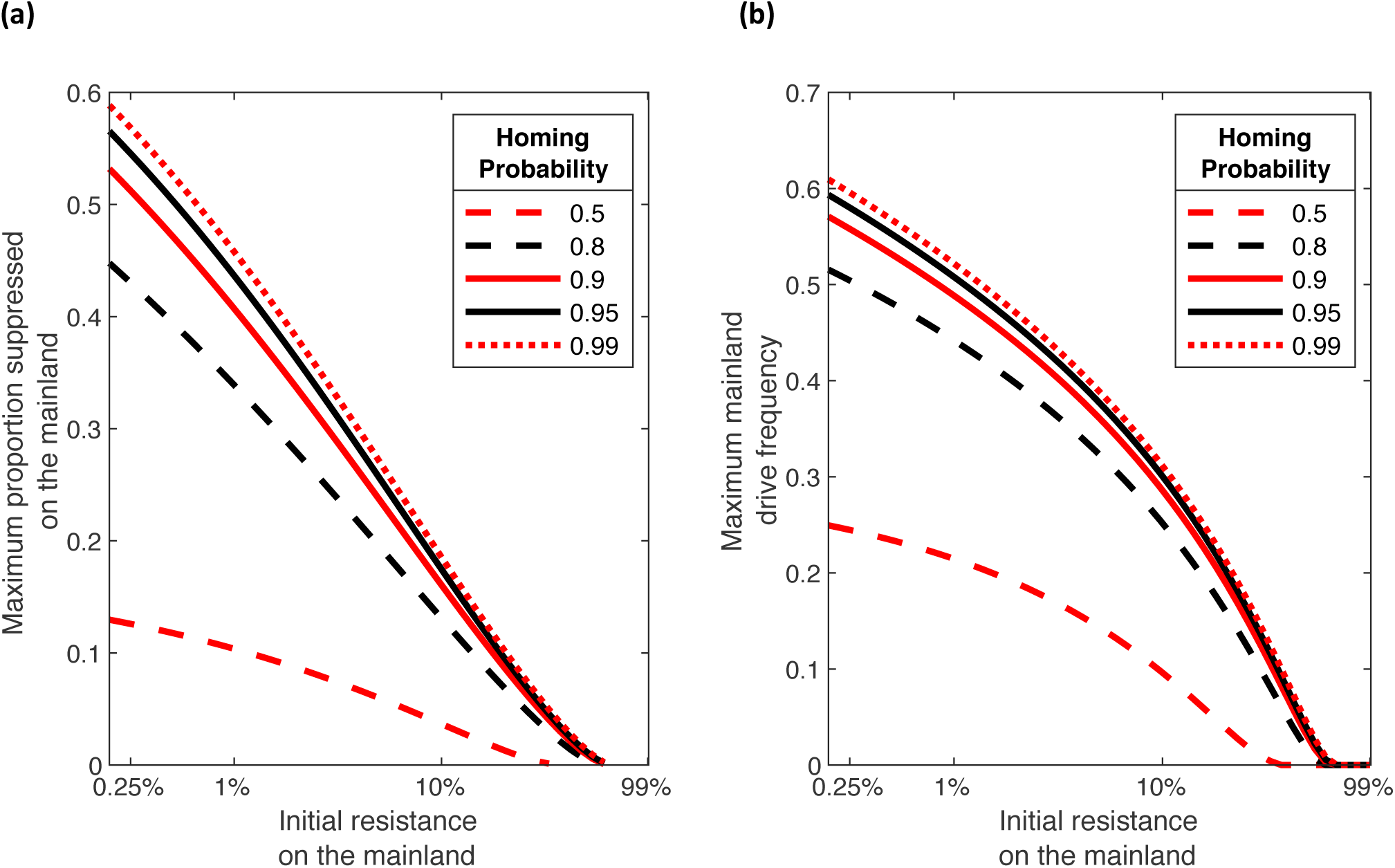
(Panel a) Maximum population suppression seen on the mainland and (Panel b) maximum drive frequency seen on mainland for various levels of mainland initial resistant allele frequency, and for different values of the homing probability. In all cases, drive parameters are taken to equal *s*=0.8 and *h*=0.3. All other parameters are as in Figure 3. of the main text.

#### S.3.2. Sensitivity to Demographic and Density Dependence Parameters

##### S.3.2.1 Logistic Model

We also explore the dependence of our results to demographic and density dependence parameters. The dynamics of the deterministic logistic model involves two parameter combinations: the per-capita growth rate of the population at low densities (*λ−ρ*) and the per-capita rate of change of the population growth rate. In order to make comparisons across parameter values, we choose to keep the equilibrium population size fixed. This leaves us with the single parameter combination *λ−ρ* that can be varied in order to explore the impact of demographic parameters within this demographic model. Here, we choose to vary *λ* in order to achieve this. The primary impact of varying parameters in this way is to change the stability of the positive equilibrium of the logistic model. A natural way to characterize this is by calculating the *return time* of the equilibrium, a measure of the time taken for perturbations of the population about its equilibrium to decay that is commonly used in the theoretical ecology literature^8^. More precisely, the return time is calculated as the reciprocal of the absolute value of the largest eigenvalue of the Jacobian matrix of the model at its positive equilibrium, 1/|*f*’(*N*\*)|, which for this model simply equals 1/(*λ−ρ*). Longer return times mean that the population responds more slowly to perturbations away from equilibrium (the equilibrium is “less stable” in this sense).

Figure S.8(a) shows how the maximum suppression seen on the mainland varies as *λ* is changed, and Figure S.8(b) reinterprets these results in terms of the resulting return time to equilibrium. We see that the maximum suppression seen on the mainland varies in an intuitive fashion as *λ* is changed. Longer equilibrium return times lead to higher levels of suppression: longer return times mean that the population responds more slowly to perturbations in its size and thus the fitness cost imposed by drive can push the population down to lower levels. The variation in the maximum suppression as *λ* changes can also be expressed in terms of the elasticity^9^, the ratio between the percentage change in maximum suppression and the percentage change in *λ*, calculated at the baseline parameter set and assuming that the percentage change in *λ* is small. This elasticity is equal to −1.22, which means that an *X*% change in *λ* leads to an approximate change of −1.22*X*% in the maximum suppression.

Figure S.8(c) shows that the maximum mainland drive frequency depends only weakly on the demographic parameter *λ*. This is not surprising: at the level of a single patch, population genetics and ecological dynamics are uncoupled (see^7^, for example). The weak dependence seen in the figure reflects the impact of migration between two populations whose sizes are varying. Changing the level of density dependence on the island leads to changing the timing of population reduction on the island, hence changing the numbers of drive individuals moving from the island to mainland over time. Similarly, changing the level of density dependence on the mainland changes the timing of population reduction on the mainland. This, in turn, changes the impact of arriving drive-bearing individuals: how they alter mainland drive frequency depends on the relative numbers of arriving and mainland individuals.

**Supplemental Figure 8:**
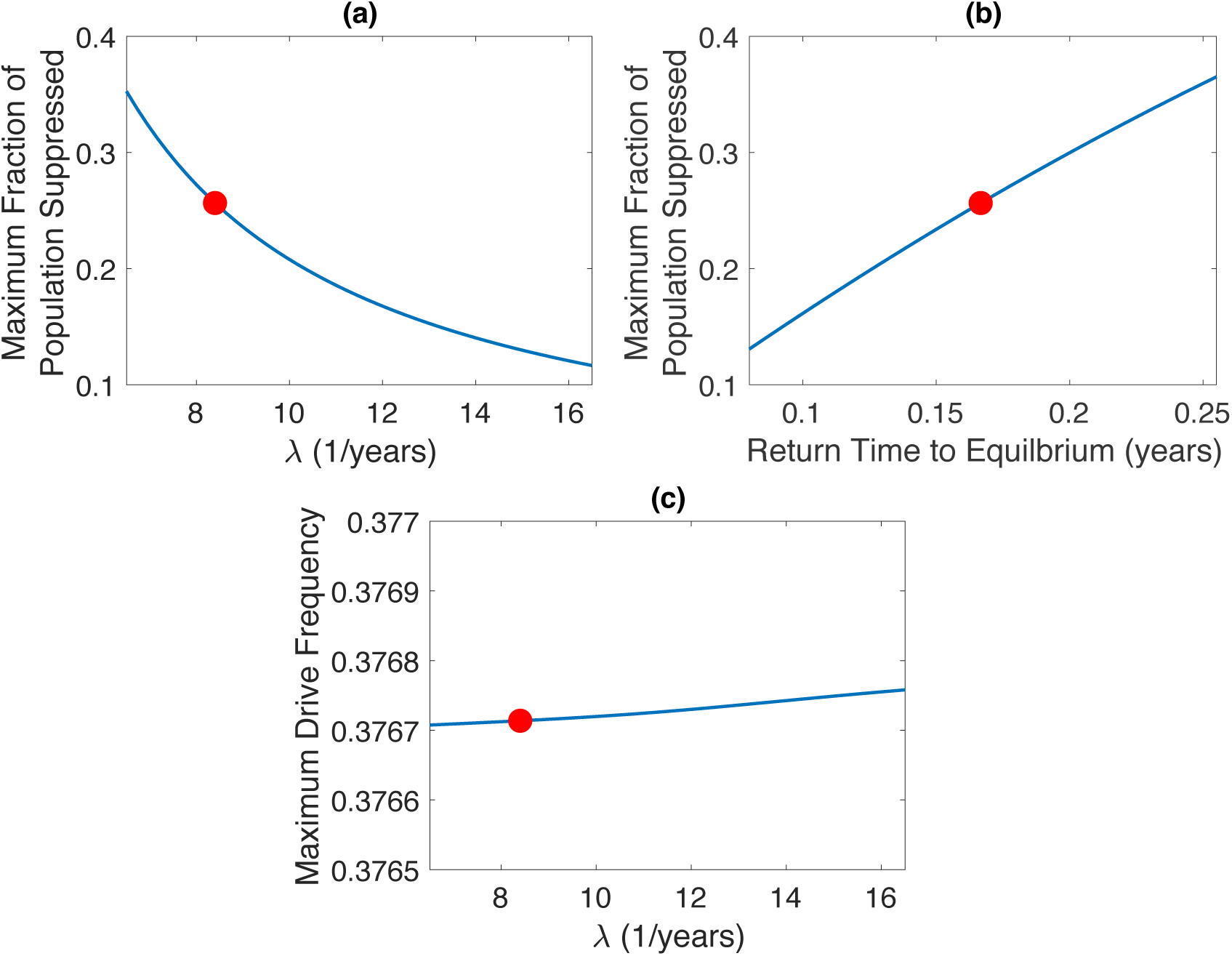
Dependence of: (panel a) the maximum suppression observed on the mainland, and (panel c) the maximum drive frequency seen on the mainland as the intrinsic per-capita growth rate of the population (*λ*) is varied. Panel (b) reinterprets the results of panel (a) in terms of the return time to equilibrium (see text for more details). All other parameters are kept fixed at the baseline values used in Figure 3, except for the parameters *q* that describe the linear decline in the per-capita birth rate with increasing population size. The *q* parameters are varied so as to keep island and mainland population sizes fixed as *λ* is changed. The red dot on each panel corresponds to the baseline set of demographic parameters used in Figure 3 of the main text.

##### S.3.2.2. Generalized Logistic Model

In the previous section, the sensitivity of model results to the demographic assumptions of the logistic model was explored. As discussed, fixing the equilibrium population size leaves us with relatively little ability to impact demography within the confines of the logistic model framework. We can make a more general exploration of the impact of density dependence by employing a slightly more general population dynamics framework, the generalized logistic model^10, 11^. Here we assume that the linearly increasing per-capita death rate of the logistic model is replaced by a nonlinear term that involves the exponent *β*-1 :

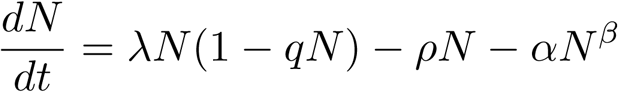

A β value of 2 corresponds to the logistic model employed in the main text and by Backus & Gross^12^. Values of β above 2 correspond to stronger density dependence, values below 2 to weaker density dependence.

As before, we keep the equilibrium population size constant when making comparisons. There are various ways to do this while varying *β*, but we employ the simplest choice: we keep the parameters *λ*, *q* and *ρ* fixed, and choose an appropriate value of *α*. To further simplify our exploration here, we only employ the generalized logistic model on the mainland, maintaining the baseline logistic dynamics on the island. (This means that the timeseries of numbers of migrants arriving on the mainland is kept the same as we vary *β*, and we explore how changing density dependence on the mainland alters the impact of these migrants on the mainland population.) Furthermore, we assume that density dependence only occurs in the death process on the mainland, i.e. we set the mainland value of *q* equal to zero.

As expected higher levels of suppression are seen for weaker density dependence (*β*<2) and lower levels for stronger density dependence (*β*>2), compared to the logistic model (*β*=2). The elasticity, calculated at the baseline level of the parameter, is −1.22.

**Supplemental Figure 9.**
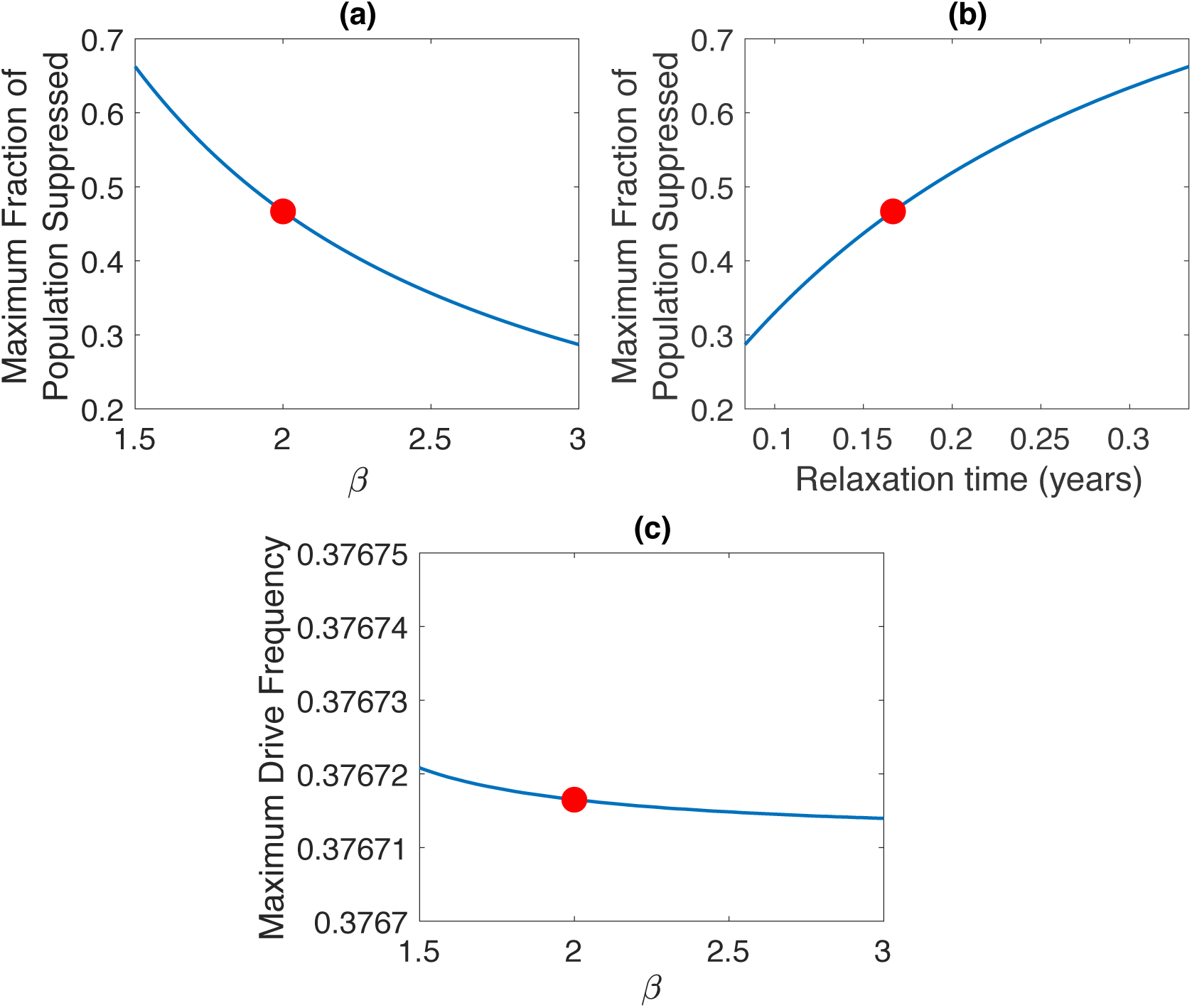
Dependence of: (panel a) the maximum suppression observed on the mainland, and (panel c) the maximum drive frequency seen on the mainland as the parameter *β* that determines the strength of density dependence is varied. Panel (b) reinterprets the results of panel (a) in terms of the return time to equilibrium (see text for more details). It is assumed that density dependence on the mainland only occurs via deaths (i.e. the mainland *q* parameter is set equal to zero). All other parameters are kept fixed at the baseline values used in Figure 3 of the main text, except for the coefficient *α* of the density dependent (nonlinear) mortality term on the mainland. This parameter is varied so as to keep the baseline (pre-release equilibrium) mainland population size fixed as *β* is changed. The red dot on each panel corresponds to the baseline set of demographic parameters used in Figure 3 of the main text.

### S.4. Stochastic Model

We formulate a stochastic model in the familiar way by reinterpreting the birth, death and migration rates of the deterministic model as rates at which discrete transitions occur in a continuous-time, discrete state Markov chain model (see, for example, Renshaw^13^). Numbers of individuals of each genotype are now integer-valued quantities and processes in the model occur discretely: for instance, migration events involve the movement of a single individual from the island to the mainland. Standard stochastic simulation methods can be used to produce a collection of realizations (simulation runs) of the model.

Because the model is stochastic, repeated simulation starting from the same initial condition leads to variation in observed dynamics. One way to summarize this variability is by a histogram that depicts the distribution of model outcomes across a collection of simulation runs.

An important difference between stochastic and deterministic models is that invasion of drive is no longer guaranteed in the stochastic model, even for choices of parameters and initial conditions for which invasion is certain in the deterministic model. For instance, just by chance it could happen that a drive individual that arrives on the mainland dies before having any offspring there. In general, branching process theory can be used to calculate the probabilities that the arrival of a single drive individual will lead to successful invasion of drive or the failure of drive to spread^14^. These probabilities naturally depend on a number of drive-related parameters. Furthermore, repeated introduction of drive is more likely to lead to successful establishment of drive than a single introduction. For the baseline drive parameters used in the main text (and in this Appendix), numerical simulation shows that the probability of successful spread of drive following the arrival of a single drive individual is approximately 0.315.

For parameters corresponding to Figure 3. in the main text, with a mainland population of *N* = 100,000, a per-capita migration rate of 0.012/year and the rather pessimistic assumption that the frequency of the resistant allele on the mainland is only 5% (target allele frequency of 95%), we see that the stochastic model gives results that correspond closely to those obtained from the deterministic model. We see a relatively small variation in both the maximum level of suppression and maximum drive frequency seen on the mainland about the values predicted by the deterministic model (Figure S.10).

For a lower level of migration, μ=0.0012/year, we see (Figure S.11) that drive fails to invade on the mainland in a large number of realizations (5,356 out of 10,000). This occurs because at this level of migration, no drive individuals migrated to the mainland before extinction happened on the island in about 14% of the realizations (1,440 out of 10,000). Even if drive-bearing individuals arrived on the mainland, invasion was not guaranteed to occur: drive failed to invade in 3,916 out of the 8,560 realizations in which drive arrived on the mainland (note that some realizations involved two or more arriving migrants). Neither of these two phenomena are captured by the deterministic model, in which migration is a continually-occurring process (minute fractions of individuals continually move from island to mainland) and in which drive can invade from arbitrarily low levels (so the arrival of a fraction of a drive-bearing individual will lead to invasion of drive). Consequently, the deterministic model is in one sense overly pessimistic about the impact of drive on the mainland, in that it predicts that drive is guaranteed to (transiently) invade the mainland. On the other hand, variation about the average behavior in the stochastic model means that the deterministic model can underestimate the impact of drive when it does invade, although we see that this variation is not so large when the mainland population is large. We note that for the realizations in which drive fails to invade, we do see a non-zero maximum suppression: this reflects the variation that a stochastic wild-type population exhibits about its carrying capacity. (Note that these values would be larger if we observed the population over a longer time interval.)

**Supplemental Figure 10.**
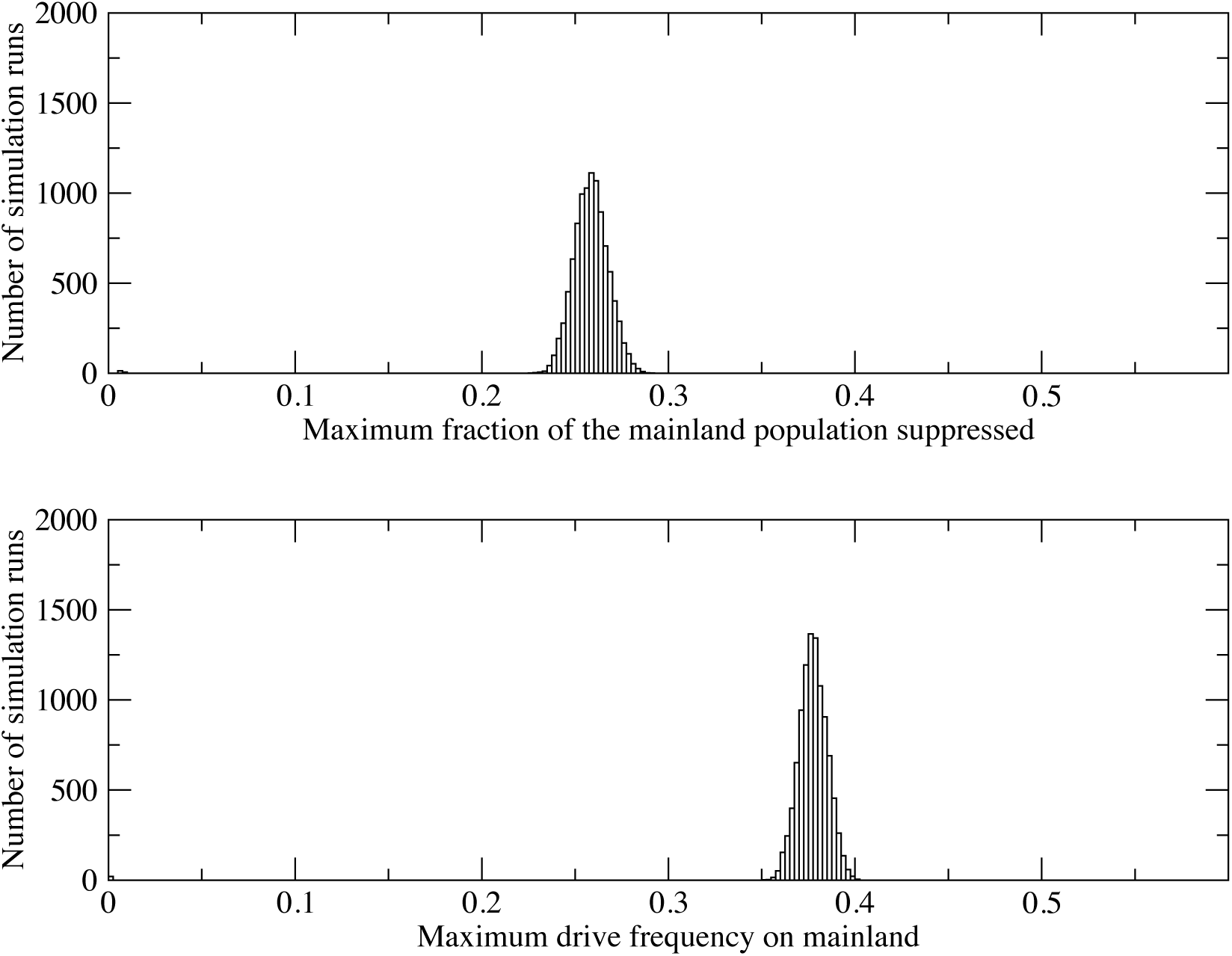
Histograms showing (a) maximum suppression seen on the mainland and (b) maximum frequency reached by drive on the mainland across 10,000 realizations of the stochastic model. Parameter values are as in Figure 3. of the main text, with a per-capita migration rate of 0.012/year and a mainland population size of 100,000.

**Supplemental Figure 11.**
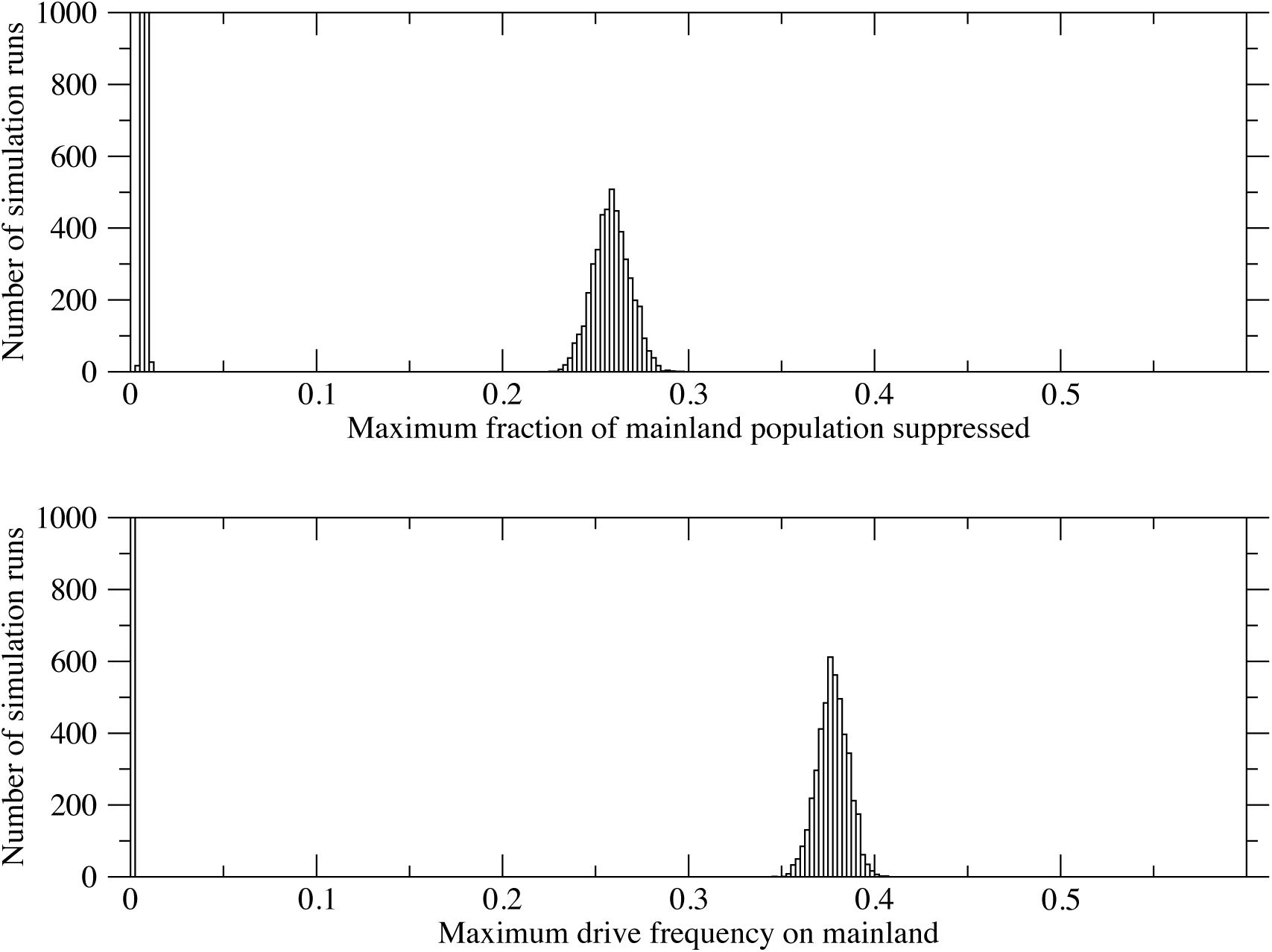
Histograms showing (a) maximum suppression seen on the mainland and (b) maximum frequency reached by drive on the mainland across 10,000 realizations of the stochastic model. Parameter values are as in Figure S.10, except that the per-capita migration rate is now lower, at 0.0012/year. Note that the observed results now exhibit bimodality: there are now a substantial number of simulation runs for which the maximum drive frequency is 0 (or low) and the maximum suppression is low. Note that the choice of scale on the vertical axis (chosen to allow the upper part of the bimodal distributions to be clearly visualized) truncates the lower parts of the bimodal distributions. 5,356 out of 10,000 simulation runs fall into the lower parts of the two distributions.

### S.5. Population Suppression and Population Replacement

We conclude with a brief description of the dynamics of the model in the case where the island population is suppressed but not eliminated. We assume that drive bears a positive fitness cost, but that the cost imposed (either at fixation or at the polymorphic equilibrium, depending on the outcome of the population genetics) is not sufficient to lead to elimination. This setting can either describe a drive that is intended to suppress the island population or one that is designed as a population replacement strategy. (Expressions for the threshold conditions governing the population genetic outcome are given in^2, 7^, as are the allele frequencies at the polymorphic equilibrium (if it exists). Alphey and Bonsall^7^ further provide threshold conditions for the elimination of a population given the population genetic outcome.)

The primary difference in the dynamics here is that drive will remain in the island population, and hence there will be continual introduction of drive to the mainland by migration. Given that drive is outcompeted on the mainland, this leads to the establishment of a polymorphic equilibrium between drive, resistant and wild-types, with the level of drive typically low. We note that if the drive fitness cost is low, these dynamics, specifically the reduction in the level of drive that occurs following its initial transient rise, can take a long time to play out.

Figure S.12 shows typical time series of the dynamics of the LFA model with a suppression drive that does not achieve elimination. Figure S.13 shows variation in the maximum level of suppression seen on the mainland, maximum mainland drive frequency, and long-term mainland drive frequency (at *t*=100 years) over a region of drive parameter space. (Note that we restrict the fitness cost *s* to be greater than 0.1 in order for our 100 year timescale to be appropriate to capture dynamics: as mentioned above, very low fitness costs lead to a very long timescale for the loss of drive on the mainland.) Figure S.14 explores the variation in the same quantities for the baseline suppression drive parameters, over a range of frequencies of resistance on the mainland and levels of migration. Results are also shown for a second set of drive parameters, with a lower fitness cost (*s*=0.3, *h*=0.3).

**Supplemental Figure 12.**
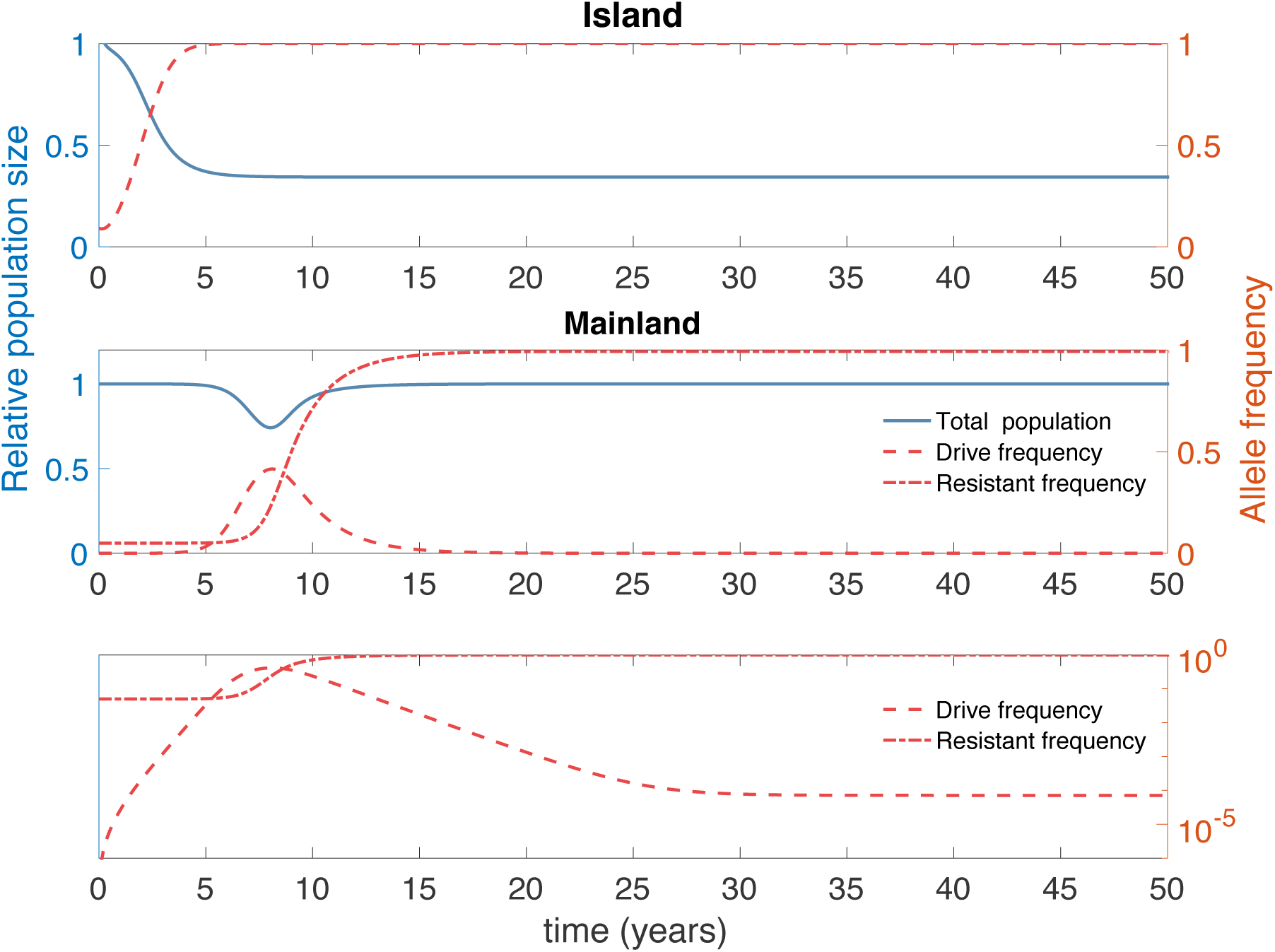
Suppression/replacement dynamics. Top Panel: Island dynamics, showing relative population size (blue solid curve; left axis) and drive allele frequency (red dashed curve; right axis). Middle Panel: Mainland dynamics: relative population size (blue solid curve; left axis), and drive and resistant allele frequencies (red dashed and red dot-dashed curves, respectively; right axis). Bottom Panel: allele frequencies as in the previous panel, but depicted on a logarithmic scale. A drive that does not achieve elimination on the island is deployed (*s*=0.6 and *h*=0.3). For these parameters, the drive achieves fixation on the island but the resulting genetic load is not sufficient to cause elimination. This leads to a continual migration of drive-bearing individuals to the mainland. Initially, dynamics on the mainland play out much as seen for an elimination drive. However, the continual reintroduction of drive to the mainland by the migrants from the island leads to a polymorphic equilibrium for which drive is present at a low level (most visible on lower panel). Other parameters are as in Figure 3 of the main text, with homing probability of 0.95, initial resistance allele frequency of 95% on the mainland and per-capita migration rate of 0.012/year.

**Supplemental Figure 13.**
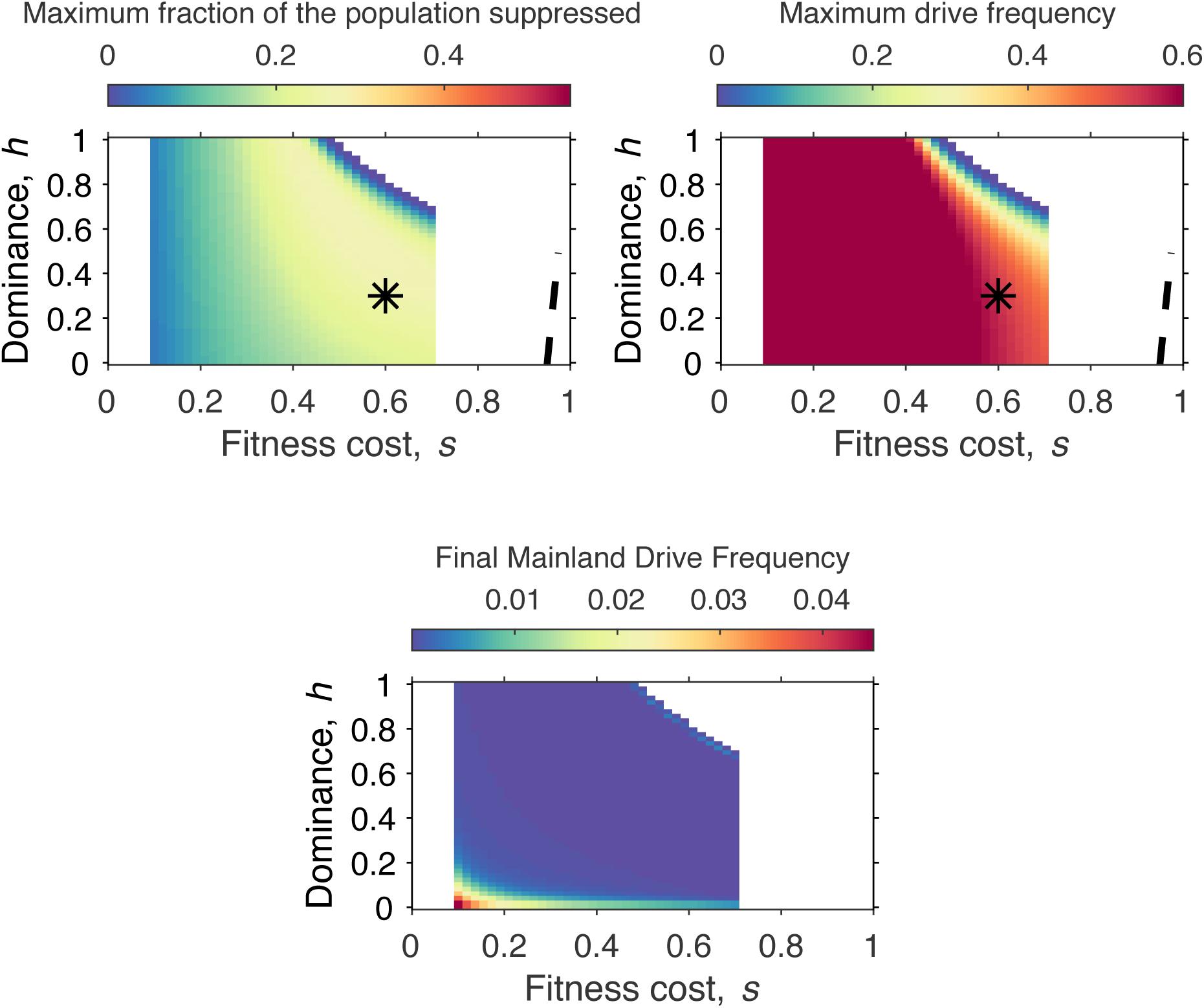
Heatmaps showing the dependence of (panel a) the maximum suppression observed on the mainland, (panel b) the maximum level of drive seen on the mainland, and (panel c) the drive frequency on the mainland after 100 years, on drive fitness parameters *s* and *h* over regions of parameter space for which the drive does not have an invasion probability and suppresses the island population, but does not does not lead to extinction. The black asterisk denotes the drive parameters used in Figures S.12. All other parameters are as in Figure S.12. The scales on the color bars in panels (a) and (b) are chosen to be the same as in Figures 4 and 5 of the main text. White regions on this figure denote combinations of drive parameters that either lead to threshold behavior, loss of drive or for which the drive leads to elimination of the island population. The dashed curve denotes the boundary between the region of parameter space where drive approaches fixation (to the left of the curve) and where drive approaches a polymorphic equilibrium with wild-type (to the right of the curve). (Note that with the homing probability of 0.95 used here, the dashed curve lies in the region of drive space that leads to extinction of the island population.)

**Supplemental Figure 14.**
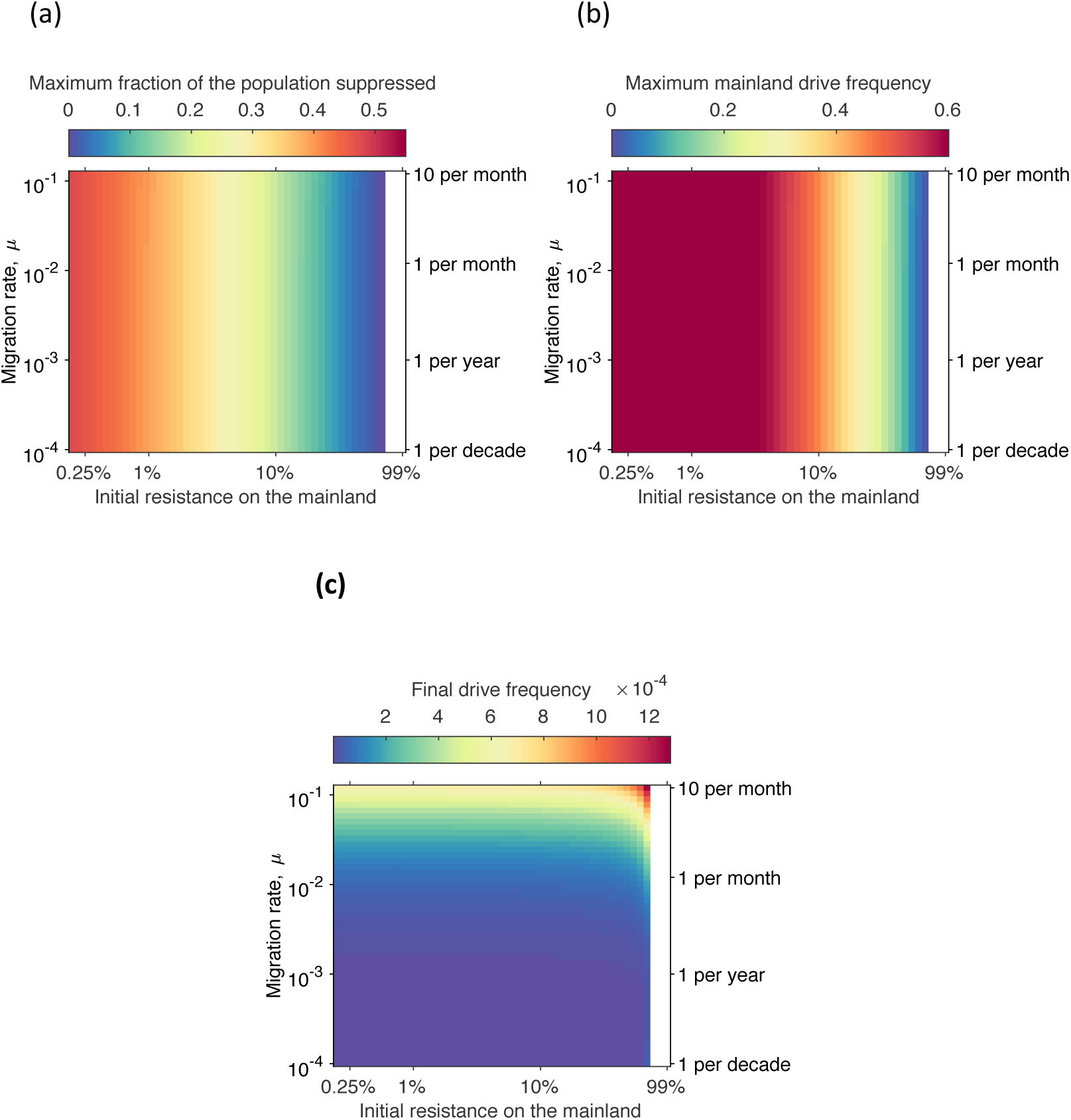
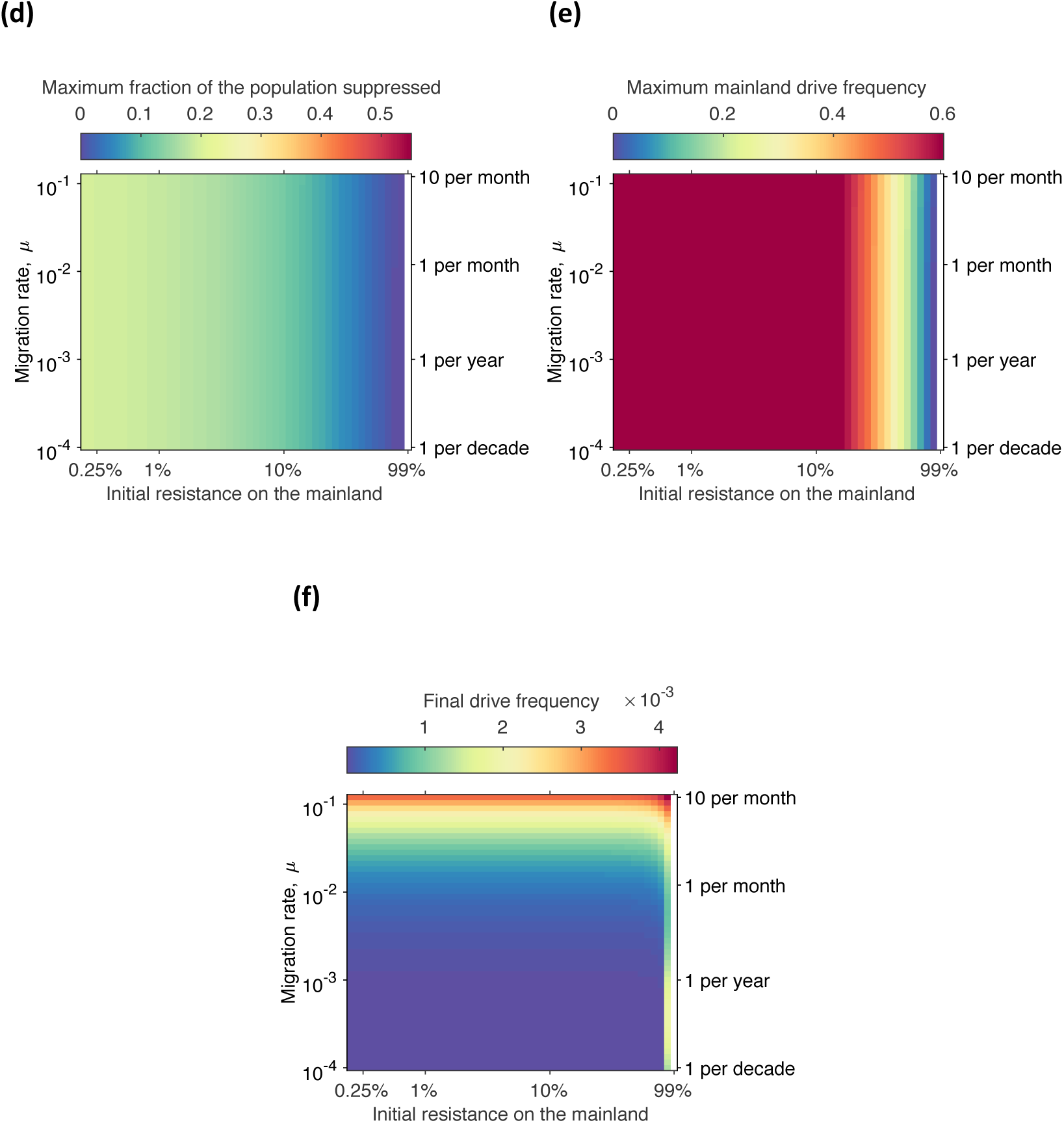
Heatmaps showing the dependence of (panel a) the maximum suppression observed on the mainland, (panel b) the maximum level of drive seen on the mainland, and (panel c) the drive frequency on the mainland after 100 years, on initial level of resistance on the mainland and the migration rate, for drive parameters *s*=0.6 and *h*=0.3 that lead to threshold-free suppression but not extinction of the island population. Panels (d), (e), and (f) show the same information but for a drive with *s*=0.3 and h=0.3. The scales on the color bars in panels (a) and (d), and (b) and (e) are chosen to be the same as in Figures 4 and 5 of the main text. All other parameters are as in Figure S.12.

## References

1. Curtis, C. F. (1968). Possible use of translocations to fix desirable genes in insect pest populations. Nature 218(5139), 368–369.

2. Burt, A. (2003). Site-specific selfish genes as tools for the control and genetic engineering of natural populations. Proc. R. Soc. Lond. B 270(1518), 921–928.

3. Gould, F. (2008). Broadening the application of evolutionarily based genetic pest management. Evolution 62(2), 500–510.

4. Esvelt, K., Smidler, A. L., Catteruccia, F., & Church, G. M. (2014). Concerning RNA-guided gene drives for the alteration of wild populations. eLIFE 3, e03401.

5. Champer, J., Buchman, A., & Akbari, O. S. (2016). Cheating evolution: engineering gene drives to manipulate the fate of wild populations. Nat. Rev. Genet. 17, 146–159.

6. Spielman, A. (1994). Why entomological antimalaria research should not focus on transgenic mosquitoes. Parasitol. Today 10,374–376.

7. Alphey, L., Beard, C. B., Billingsley, P., Coetzee, M., Crisanti, A., Curtis, C., Eggleston, P., Godfray, C., Hemingway, J., Jacobs-Lorena, M., James, A. A., Kafatos, F. C., Mukwaya, L. G., Paton, M., Powell, J. R., Schneider, W., Scott, T. W., Sina, B., Sinden, R., Sinkins, S., Spielman, A., Toure, Y., & Collins, F. H. (2002). Malaria control with genetically manipulated insect vectors. Science 298, 119–121.

8. Macer, D. (2005). Ethical, legal and social issues of genetically modifying insect vectors for public health. Insect Biotech. Mol. Biol. 35, 649–660.

9. Marshall, J. M. (2010). The Cartagena Protocol and genetically modified mosquitoes. Nature Biotech. 28, 896–897.

10. Akbari, O. S., Bellen, H. J., Bier, E., Bullock, S. L., Burt, A., Church, G. M., Cook, K. R., Duchek, P., Edwards, O. R., Esvelt, K. M., Gantz, V. M., Golic, K. G., Gratz, S. J., Harrison, M. M., Hayes, K. R., James, A. A., Kaufman, T. C., Knoblich, J., Malik, H. S., Matthews, K. A., O’Connor-Giles, K. M., Parks, A. L., Perrimon, N., Port, F., Russell, S., Ueda, R., & Wildonger, J. (2015). Safeguarding gene drive experiments in the laboratory. Science, 349, 927–929.

11. National Academies of Sciences, Engineering, and Medicine (2016). Gene Drives on the Horizon: Advancing Science, Navigating Uncertainty, and Aligning Research with Public Values. Washington, DC: The National Academies Press. doi: 10.17226/23405.

12. Esvelt, K. M., & Gemmell, N. J. (2017). Conservation demands safe gene drive. PLoS Biology 15(11), e2003850.

13. Marshall, J. M. & Akbari, O. S. (2018). Can CRISPR-Based Gene Drive Be Confined in the Wild? A Question for Molecular and Population Biology. ACS Chem. Biol. 13(2), 424–430.

14. Davis, S., Bax, N., & Grewe, P. (2001). Engineered underdominance allows efficient and economical introgression of traits into pest populations. J. Theor. Biol. 212(1), 83–98.

15. Altrock, P. M., Traulsen, A., Reeves, R. G., & Reed, F. A. (2010). Using underdominance to bi-stably transform local populations. J. Theor. Biol. 267, 62–75.

16. Barton, N. H. (1979). The dynamics of hybrid zones. Heredity 43, 341–359.

17. Barton, N. H., & Turelli, M. (2011). Spatial waves of advance with bistable dynamics: Cytoplasmic and genetic analogues of Allee effects. Am. Nat. 178(3), E48–E75.

18. Gould, F., Huang, Y., Legros, M., & Lloyd, A. L. (2008). A Killer–Rescue system for self-limiting gene drive of anti-pathogen constructs. Proc. R. Soc. Lond. B 275(1653), 2823–2829.

19. Rasgon, J. L. (2009). Multi-locus assortment (MLA) for transgene dispersal and elimination in mosquito populations. PLoS ONE, 4, e5833.

20. Burt, A., & Deredec, A. (2018). Self-limiting population genetic control with sex-linked genome editors. Proc. R. Soc. Lond. B, 285, 20180776.

21. Noble, C., Min, J., Olejarz, J., Buchthal, J., Chavez, A., Smidler, A., DeBenedictis, E., Church, G., Nowak, M., & Esvelt, K. (2016). Daisy-chain gene drives for the alteration of local populations. bioRxiv 057307. doi:10.1101/057307.

22. Dhole, S., Vella, M. R., Lloyd, A. L., & Gould, F. (2018). Invasion and migration of spatially self-limiting gene drives: A comparative analysis. Evol. Applic. 11(5), 794–808.

23. Alphey, N., & Bonsall, M. B. (2014). Interplay of population genetics and dynamics in the genetic control of mosquitoes. J. R. Soc. Interface, 11(93), 20131071.

24. Khamis, D., El Mouden, C., Kura, K., & Bonsall, M. B. (2018). Ecological effects on underdominance threshold drives for vector control. J. Theor. Biol. 456, 1–15.

25. Kimura, M., & Ohta, T. (1968). The average number of generations until fixation of a mutant gene in a finite population. Genetics 61, 763–771.

26. Frankham, R. (1997). Do island populations have lower genetic variation than mainland populations? Heredity 78, 311–327.

27. Hsu, P. D., Scott, D. A., Weinstein, J. A., Ran, F. A., Konermann, S., Agarwala, V., Li, Y., Fine, E. J., Wu, X., Shalem, O., Cradick, T. J., Marraffini, L. A., Bao, G., & Zhang, F. (2013). DNA targeting specificity of RNA-guided Cas9 nucleases. Nature Biotech. 31, 827–832.

28. Howald, G., Donlan, C. J., Galvan, J. P., Russell, J. C., Parkes, J., Samaniego, A., Wang, Y., Veitch, D., Genovesi, P., Pascal, M., Saunders, A., & Tershy, B. (2007). Invasive rodent eradication on islands. Conserv. Biol. 21(5), 1258–1268.

29. Cuthbert, R., & Hilton, G. (2004). Introduced house mice *Mus musculus*: a significant predator of threatened and endemic birds on Gough Island, South Atlantic Ocean? Biol. Cons. 117, 483–489.

30. Towns, D. R., Atkinson, I. A. E., & Daughtery, C. H. (2006). Have the harmful effects of introduced rats on islands been exaggerated? Biol. Invasions 8, 863–891.

31. Campbell, K. J., Beek, J., Eason, C. T., Glen, A. S., Godwin, J., Gould, F., Holmes, N. D., Howald, G. R., Francine M. Madden, F. M., Ponder, J. B., Threadgill, D. W., Wegmann, A. S., & Baxter, G. S. (2015) The next generation of rodent eradications: Innovative technologies and tools to improve species specificity and increase their feasibility on islands. Biol. Cons. 185, 47–58.

32. Backus, G. A., & Gross, K. (2016). Genetic engineering to eradicate invasive mice on islands: modeling the efficiency and ecological impacts. Ecosphere 7(12), e01589.

33. Robert, M. A., Okamoto, K., Lloyd, A. L., & Gould, F. (2013). A reduce and replace strategy for suppressing vector-borne diseases: insights from a deterministic model. PLoS One 8(9), e73233.

34. Prowse, T. A. A., Cassey, P., Ross, J. V., Pfitzner, C., Wittmann, T. A., & Thomas, P. (2017) Dodging silver bullets: good CRISPR gene-drive design is critical for eradicating exotic vertebrates. Proc. R. Soc. Lond. B 284, 20170799.

35. Nathan, H. W., Clout, M. N., MacKay, J. W., Murphy, E. C., & Russell, J. C. (2015). Experimental island invasion of house mice. Popul. Ecol. 57(2), 363–371.

36. Deredec, A., Burt, A., & Godfray, C. (2008). The population genetics of using homing endonuclease genes (HEGs) in vector and pest management. Genetics 179, 2013–2026.

37. Unckless, R. L., Messer, P. W., Connallon, T., & Clark, A. G. (2015). Modeling the manipulation of natural populations by the mutagenic chain reaction. Genetics 201(4), 425–531.

38. Magori, K., & Gould, F. (2006). Genetically engineered underdominance for manipulation of pest populations: A deterministic model. Genetics 172, 2613–2620.

39. Dhole, S., Lloyd, A. L., & Gould, F. (2019). Tethered homing gene drives: A new design for spatially restricted population replacement and suppression. Evol. Applic. 12(8), 1688–1702.

40. Wilkins, K. E., Prowse, T. A. A., Cassey, P., Thomas, P. Q., & Ross J. V. (2018) Pest demography critically determines the viability of synthetic gene drives for population control. Math. Biosci. 305, 160–169.

41. Browne, R A.1977. Genetic variation in island and mainland populations of *Peromyscus leucops*. Am Midl. Nat, 97, 1–9.

42. White, T. A., & Searle, J. B. (2007). Genetic diversity and population size: island populations of the common shrew, *Sorex araneus*. Mol. Ecol. 16, 2005–2016.

43. Harradine, E., How, R. A., Schmitt, L. H., & Spencer, P. B. S. (2015). Island size and remoteness have major conservation significance for how spatial diversity is partitioned in skinks. Biodiv. Cons. 24, 2011–2029.

44. Champer, J., Liu, J., Oh, S. Y., Reeves, R., Luthra, A., Oakes, N., Clark, A. G., & Messer, P. W. (2018). Reducing resistance allele formation in CRISPR gene drive. Proc. Natl. Acad. Sci. USA 115, 5522–5527.

45. Piaggio, A. J., Segelbacher, G., Seddon, P. J., Alphey, L., Bennett, E. L., Carlson, R. H., Friedman, R. M., Kanavy, D., Phelan, R., Redford, K. H., Rosales, M., Slobodian, L., & Wheeler, K. (2017) Is it time for synthetic biodiversity conservation? Trends Ecol. Evol. 32, 97–107.

## References

1. Strogatz, S. H. (2014) Nonlinear Dynamics and Chaos, Second Edition. Westview Press.

2. Deredec, A., Burt, A., & Godfray, C. (2008). The population genetics of using homing endonuclease genes (HEGs) in vector and pest management. Genetics 179, 2013–2026.

3. Vella, M. R., Gunning, C. E., Lloyd, A. L., & Gould, F. (2017) Evaluating strategies for reversing CRISPR-Cas9 gene drives. Scientific Reports 7, 11038.

4. Prowse, T. A. A., Adikusuma, F., Cassey, P., Thomas, P., & Ross, J. V. (2019). A Y-chromosome shredding gene drive for controlling pest vertebrate populations. eLife 8, e41873.

5. Saltelli, A., Ratto, M., Andres, T., Campolongo, F., Cariboni, J., Gatelli, G., Saisana, M., & Tarantola, S. (2008). Global Sensitivity Analysis, the Primer. Wiley.

6. Pianosi, F., Sarrazin, F., & Wagener, T. (2015). A Matlab toolbox for Global Sensitivity Analysis. Environ. Model. Software 70, 80–85.

7. Alphey, N., & Bonsall, M. B. (2014). Interplay of population genetics and dynamics in the genetic control of mosquitoes. J. R. Soc. Interface, 11(93), 20131071.

8. DeAngelis, D. L., & Waterhouse, J. C. (1987). Equilibrium and Nonequilibrium Concepts in Ecological Models. Ecol. Monogr. 57, 1–21.

9. Benton, T. G., & Grant, A. (1999). Elasticity analysis as an important tool in evolutionary and population ecology. Trends Ecol. Evol. 14, 467–471.

10. Robert, M. A., Okamoto, K., Lloyd, A. L., & Gould, F. (2013). A reduce and replace strategy for suppressing vector-borne diseases: insights from a deterministic model. PLoS One 8(9), e73233.

11. Prowse, T. A. A., Cassey, P., Ross, J. V., Pfitzner, C., Wittmann, T. A., & Thomas, P. (2017) Dodging silver bullets: good CRISPR gene-drive design is critical for eradicating exotic vertebrates. Proc. R. Soc. Lond. B 284, 20170799.

12. Backus, G. A., & Gross, K. (2016). Genetic engineering to eradicate invasive mice on islands: modeling the efficiency and ecological impacts. Ecosphere 7(12), e01589.

13. Renshaw, E. (1991). Modelling Biological Populations in Space and Time (Cambridge Studies in Mathematical Biology). Cambridge: Cambridge University Press.

14. Marshall, J.M. (2009). The effect of gene drive on containment of transgenic mosquitoes. J. Theor. Biol. 258, 250–265.

